# The Placental Transcriptome Serves as a Mechanistic Link between Prenatal Phthalate Exposure and Placental Efficiency

**DOI:** 10.1101/2025.08.05.664214

**Authors:** Mariana Parenti, Samantha Lapehn, James MacDonald, Theo Bammler, Adam Szpiro, Marnie Hazlehurst, Drew B. Day, Ciara Thoreson, Kurunthachalam Kannan, Nicole R. Bush, Kaja Z. LeWinn, Qi Zhao, Sheela Sathyanarayana, Alison G. Paquette

## Abstract

Prenatal exposure to phthalates, pervasive endocrine-disrupting chemicals, has been linked to child health outcomes, including prematurity and low birthweight. Placental transcriptomics data can reveal mechanisms by which environmental toxicants alter placental and fetal growth. This study aims to investigate the placental transcriptome as a mediator between prenatal maternal urinary phthalate metabolites and placental efficiency. We identified significant associations between maternal urinary concentrations of two phthalate metabolites and the placental transcriptome (132 genes and 27 gene modules). 7 genes and 3 gene modules exhibited significant consistent mediation of the relationship between phthalates and placental efficiency measures. These genes were involved in syncytialization, metabolism, DNA damage and cellular senescence, and steroid biosynthesis—processes essential to fetal growth and development because of the placenta’s role in nutrient supply, hormone production, and detoxification. These findings suggest a key mediating role of the placental transcriptome in toxicological mechanisms by which phthalates may disrupt fetal growth.

**Teaser:** Placental gene expression mediates the relationship between prenatal phthalate exposure and fetal growth measures.

## Introduction

Phthalates are plasticizers present in numerous consumer products, leading to ubiquitous human exposure (*1*, *2*). Phthalates are hydrolyzed to mono-esters and transformed into secondary metabolites based on their chemical structure and molecular weight, resulting in a variety of metabolites which the fetus and placenta are exposed to during pregnancy, and are detectable in urine (*3*). Increased prenatal phthalate exposure is associated with impaired fetal (*4–6*) and childhood growth including weight and BMI (*7–12*). These studies suggest that phthalate exposure during pregnancy may lead to disruptions to in the *in utero* environment that have lasting effects on developing children from infancy into childhood.

Phthalates cross the placental barrier and interact with the fetus to alter programming of mechanisms involved in development and growth (*13*). The placenta plays a central role in the interplay between the maternal and fetal environments and early life programming (*14*, *15*) as it regulates fetal nutrient transport, oxygen and waste exchange, and endocrine signaling (*14*, *16*). Phthalates have several established mechanisms of toxicity that may disrupt placental growth and angiogenesis, ultimately leading to impaired fetal growth. Phthalates alter placental structural formation by disrupting trophoblast differentiation (*18*), inhibiting trophoblast invasion (*19*), and causing structural abnormalities (*20*). Phthalates also disrupt placental angiogenesis (*21*, *22*), which can impair the placenta’s capacity to effectively transport nutrients needed for growth (*23*). Phthalates activate the aryl hydrocarbon receptor, leading to initiation of pro-apoptotic pathways (*24*, *25*). Placental cellular apoptosis can impair trophoblast function and reduce nutrient transport, and has been noted in intra-uterine growth restriction (*26*). Phthalates also activate Peroxisome Proliferator Activated Receptor Gamma (PPARG) (*25*, *27*, *28*). Placental PPARG expression is positively associated with both placental and fetal weight (*29*). These experimental studies demonstrate potential mechanisms by which phthalates could disrupt fetal and child growth, but the specific mechanisms have not been established.

Placental weight is predictive of birthweight, explaining up to 32% of the variability of birth weight (*17*). Placental efficiency is quantified by the birthweight:placental weight ratio (BW:PW); a proxy measure of how the placenta has adapted to meet fetal nutritional requirements, which is highly relevant to fetal growth (*30*). Both small and large placentas can be inefficient: a small placenta might contribute to lower birthweight through reduced surface area thus reducing nutrient transport to the fetus. On the other hand, the placenta is a highly metabolic organ, so a large placenta might be a nutrient sink at the expense of fetal growth (*31*). Decreased BW:PW, or a relatively large placenta compared to birthweight, is associated with increased fetal growth restriction (*30*), NICU admission (*32*), decreased APGAR scores (*32*), and preeclampsia (*33*), as well as histological changes including increased syncytial knots and intervillous fibrin (*34*). Conversely, increased BW:PW, or a relatively small placenta for a given birthweight, is associated with gestational diabetes (*33*). Placental weight is predictive of altered childhood growth (*35*), and may even be linked to later life health outcomes. For example, a Norwegian study found that infants in the highest tertile for placental inefficiency had 38% increased risk of cardiovascular disease mortality 45 years later compared to participants in the lowest tertile (*36*). Phthalate metabolites are associated with decreased placental efficiency, reduced placental vascular resistance, and lower placental weight and BW:PW ratio (*37*, *38*). The mechanisms by which phthalates disrupt placental development remain unclear.

The generation and analysis of -omics data from the placenta can reveal the effects of maternal environmental exposures on placental function and fetal development, and link environmental exposures and infant health outcomes (*39–41*). We previously generated transcriptomic signatures of prenatal phthalate exposure in the Conditions Affecting Neurocognitive Development and Learning in Early childhood (CANDLE) study, a large and diverse birth cohort from Memphis, Tennessee (*42*). The goal of this study was to expand these signatures of prenatal phthalate exposure to BW:PW ratio, placental weight (PW), and birthweight adjusted for placental weight (BW_adj_) as markers of placental efficiency. We hypothesize that phthalates disrupt placental function leading to impaired placental efficiency, and these perturbations are reflected in the placental transcriptome. We previously identified sex-specific differences in responses to other environmental chemicals, so we further considered fetal sex as a potential modifier of the relationship between phthalate metabolites and the placental transcriptome. These findings are expected to provide novel insight into how prenatal exposure to phthalates may disrupt placental function and therefore fetal growth and development.

## Results

### 1. Association between prenatal phthalate exposures and placental efficiency in PWG participants

Placental samples were biobanked and RNA sequencing was conducted for 465 participants (Ages 16-45, median age 31) enrolled in GAPPS between 2011 and 2017 by the ECHO PATHWAYS and PWG cohort studies (**Figure S1**). 212 participants (45.6%) were recruited from the Seattle, WA site and 253 participants (54.4%) were collected from the Yakima, WA site. As described in **Table S1**, the majority of participants in this cohort self-identified their race as White (*N*=358, 82.3%) and 9 participants identified as Black (2.1%), 24 participants identified as Asian (5.5%), and 44 participants identified as American Indian or Alaska Native, multiple race, or self-reported other race (10.1%), and 73 participants (16.0%) identified as Hispanic. Most participants underwent labor (88.8%) and delivered vaginally (67.7%). Urine was collected from a subset of 222 PWG participants at clinic visits during the first trimester (median visit timepoint 11.7 weeks, range 6.1-13.9 weeks), second trimester (median visit timepoint 21.9 weeks, range 14-28 weeks), and/or third trimester (median visit timepoint 33.4 weeks, range 28-41 weeks **(Figure S2A,B**), which was used for quantification of phthalate metabolites. Fifteen phthalate metabolites were detected in at least 70% of participants and di-2-ethylhexy phthalate (DEHP) was calculated as a molar sum from 5 of its metabolites. Monoethyl phthalate (MEP) had the highest median concentration (37.76 ng/ml), while mono-(7-carboxy-n-heptyl) phthalate (MCHPP) had the lowest median concentration (0.31 ng/ml) (**Figure S2C, Table S2).** All metabolite pairings had significant (p<0.05), positive Pearson correlation coefficients (**Figure S2D**). Some of the strongest correlations were among metabolites that share a parent compound including 5 metabolites of DEHP.

Placental weight and birthweight were collected through medical record abstraction from a subset of 253 PWG participants (**Table 1**). Placental efficiency was calculated as BW/PW, or how many grams of fetus were produced per gram of placenta (*43*). We observed a nonlinear relationship between placental weight and birthweight in this subset of PWG (**Figure S3A**). Using a natural cubic spline with 3 knots, we observed that the relationship was nonlinear (effective degrees of freedom=2.88, *P*<2.2×10^-16^) in models adjusted for maternal age, pre-pregnancy BMI, maternal race and ethnicity, education, smoking status, parity, gestational diabetes, hypertensive disorders of pregnancy, and fetal sex (**Figure S3B**) (*44*). The median BW:PW ratio was 7.36, with a range of 2.09-19.50. Two phthalate metabolites (MCIOP, β: - 89.68 and MCMHP, β: -103.27) were negatively associated (p<0.05) with BW_adj_ while three others (MIBP, β: 95.65; MBZP, β: 88.85; MEHP, β: 59.52) were positively associated with BW_adj_ **(Figure S4**) after adjusting for covariates (**Figure S5A**). Two phthalate metabolites were positively associated with BW:PW (MCPP, β: -0.28 and MCIOP, β: -0.24) (**Figure S4**) after adjusting for covariates (**Figure S5A**). For both analyses, there was a phthalate metabolite provisionally associated (p<0.1) with BW_adj_ (MCPP) or BW:PW (MCMHP) (**Figure S4**).

**Table 1:**
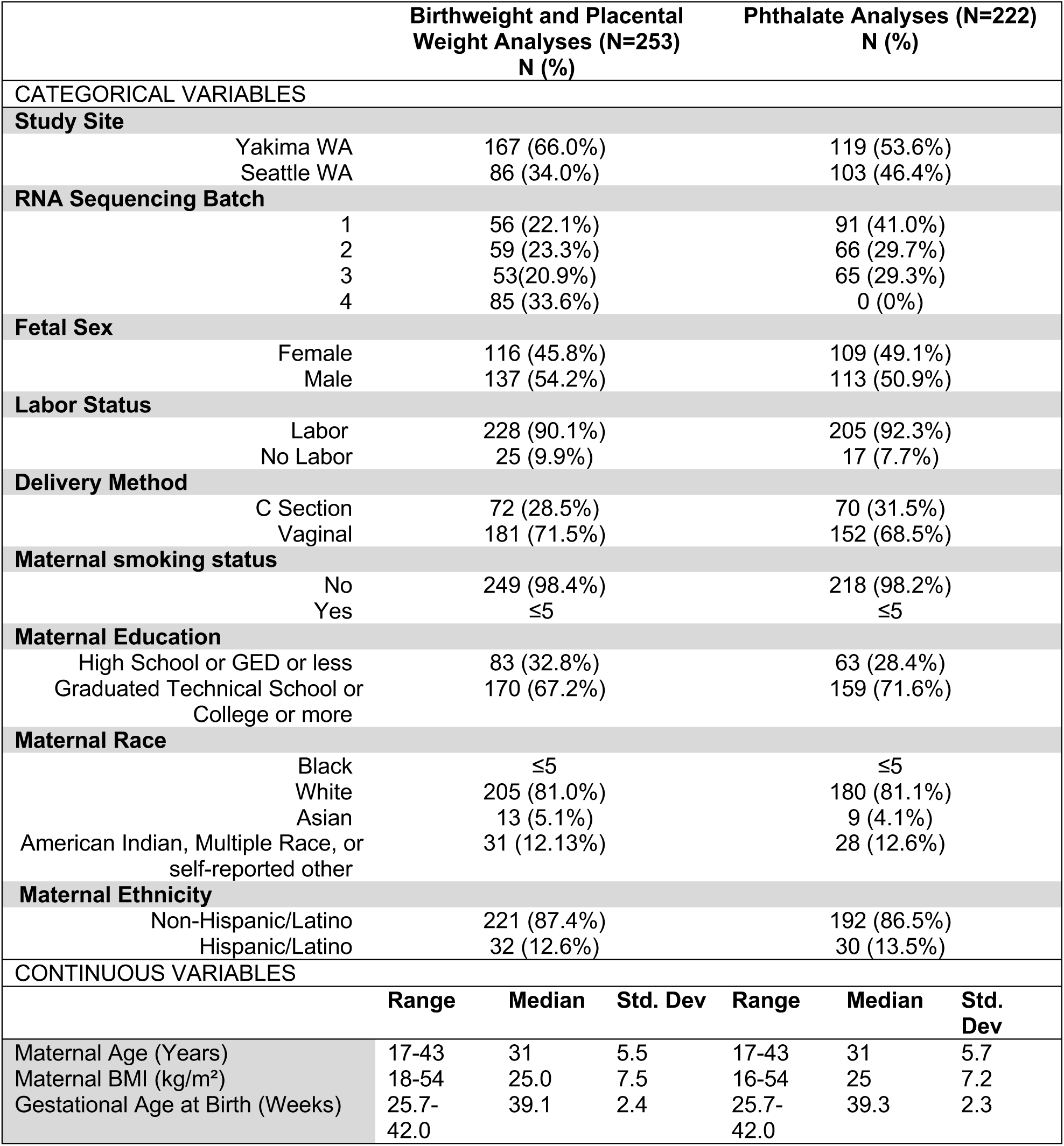
Continuous and categorical information about GAPPS cohort participants with birthweight and placental weight data included in this analysis (*N*=253) or the phthalate analyses (N=222) recruited from Seattle and Yakima, WA.

### 2. Identifying transcriptomic signatures of prenatal phthalate exposure

We generated a series of linear regression models to identify genes whose placental expression was associated with phthalate metabolites that were adjusted for covariates as described in **Figure S5B**. Two phthalate metabolites (monocarboxy isononyl phthalate (MCINP) and monomethyl phthalate (MMP)) were associated with a total of 132 genes at FDR<0.05 (**Table S3, Figure 1**). MCINP was associated with 126 genes, with the majority being positive associations, while MMP was positively associated with 6 genes. For MCINP, *FEZ1* was the most significant positively associated gene and *SPG21* was the most significant negatively associated gene. There were no genes that overlapped between the two metabolites. One gene negatively associated with MCINP, *MT-ND5*, was previously positively associated with monocarboxy isooctyl phthalate (MCIOP) in our previous analysis of prenatal phthalate exposure and the placental transcriptome in the CANDLE cohort (*42*). 20 genes were provisionally associated (p<0.005) with the phthalate mixture, including 14 genes that were positively associated and 6 genes that were negatively associated (**Table S4**). Two genes positively associated with the phthalate metabolite mixture (*RGS4* and *CEP63*) were also associated with MCINP and MMP, respectively, in the individual metabolite analysis. In both cases, the contribution of MCINP or MMP to the overall mixture effect were in the same direction as the association of the gene with the individual metabolite (**Figure S6**, **Table S4**). Across all 20 genes associated with the phthalate mixture, MECPP negatively contributed to the overall mixture effect for 15 genes, while MCINP and MIBP were the most common positive contributors to the overall mixture effects for each of the 15 genes (**Figure S6**).

**Figure 1:**
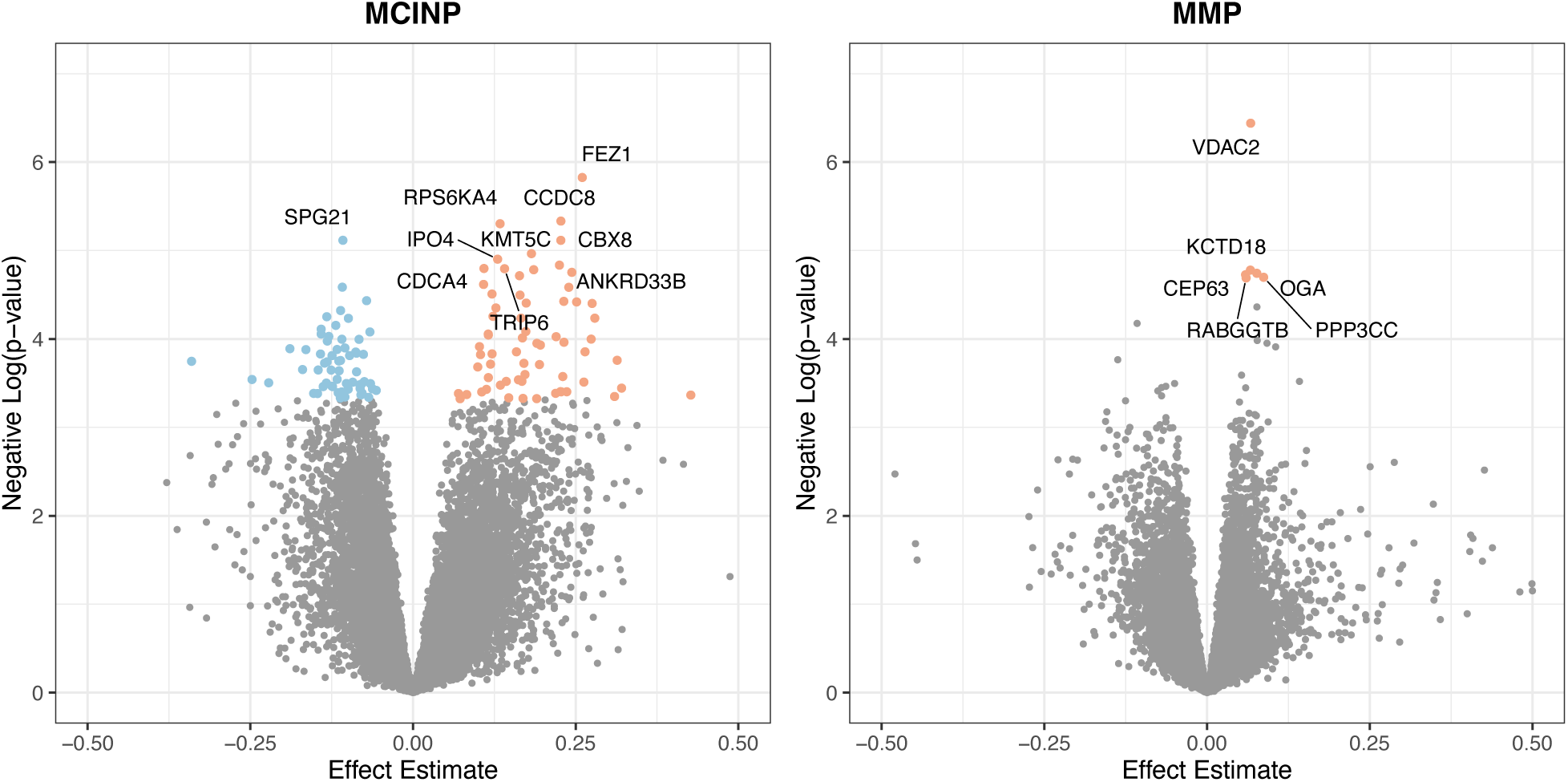
Results of differential expression analysis for phthalate metabolites that are associated with at least one gene (FDR<0.05). The 10 most significant genes based on FDR-value are labeled.

We also conducted sex-stratified analyses to investigate sex-specific effects of phthalates on the placental transcriptome. In the male stratified analysis (N=113), *DARS1* was positively associated (FDR<0.05) with MMP, and 6 genes were provisionally positively associated (p<0.005) with the phthalate metabolite mixture (**Table S4**). None of these genes were associated with individual phthalate compounds in males. Within the mixture, MBP and MECPP were exclusively positive weights for the 6 genes, while MCMHP, MEHP, and MIBP were exclusive negative weights (**Table S4**). In the female stratified model, we identified no associations between individual phthalate compounds and genes, but there were 14 provisionally associated (p<0.005) genes with the phthalate metabolite mixture. The metabolites with the most frequent negative contributions to the overall mixture effect were MCMHP and MECPP whereas MIBP and MCHHP were the most frequent metabolites that positively contributed to the overall mixture effect in females (**Table S4**).

### 3. Transcriptomic signatures of placental efficiency

To investigate placental transcriptomic signatures of placental efficiency, we evaluated the association between placental gene expression at delivery and PW, BW:PW, and BW_adj_ in models adjusted for covariates (**Figure S5C,D**). We identified 4 upregulated and 5 downregulated differentially expressed genes (DEGs) in association with BW:PW (FDR<0.05, **Table S5; Figure 2**). Sex differences in fetal growth and BW:PW have been reported (*45*, *46*), so we performed sex-stratified analyses (**Table S6**). In these models, we identified 32 DEGs associated with BW:PW in males, but no DEGs associated with BW:PW in females (**Figure S7**, FDR<0.05). *ASB2* had the strongest negative association with BW:PW in both the overall (β=-0.393) and male-specific (β=-0.495) analyses.

**Figure 2:**
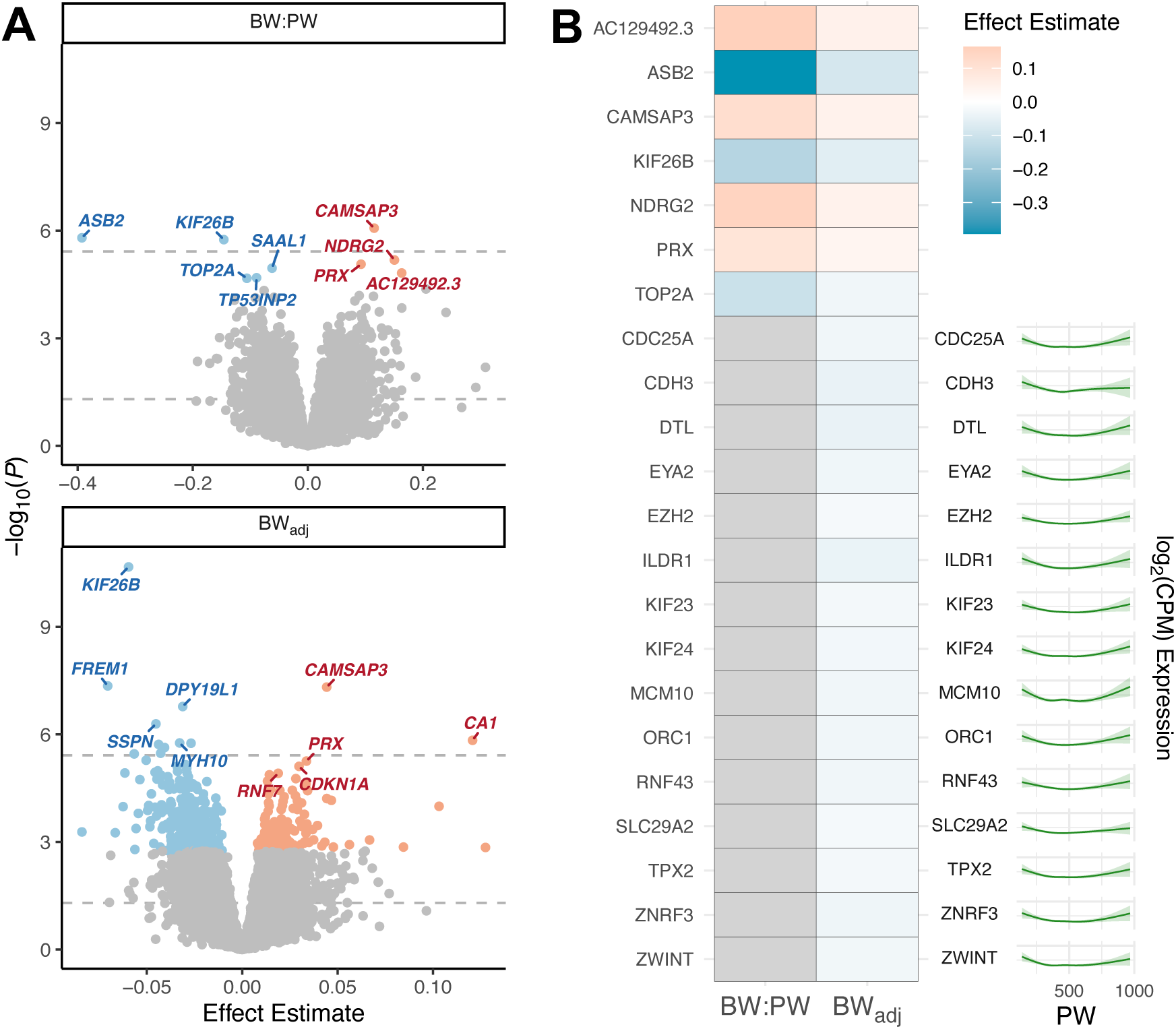
Differentially expressed genes (DEGs) associated with placental efficiency. β coefficients are presented for each 1 unit increase in the birthweight:placental weight ratio (BW:PW) and each 100-g increase in birthweight adjusted for placental weight (BW_adj_). (**A.**) Volcano plots for BW:PW and BW_adj_ showing upregulated (red) and downregulated (blue) genes meeting the FDR<0.05 threshold. The top 5 up- and downregulated genes are labeled. Dotted lines represent the Bonferroni and α=0.05 cutoffs. (**B.**) All DEGs associated with at least 2 outcomes (BW:PW, BW_adj_, or placental weight (PW)). PW had a nonlinear association with gene expression, the association between PW (g) and expression is visualized using a natural cubic spline.

Since we observed a nonlinear relationship between birthweight and placental weight in PWG participants (**Figure S3**), we hypothesized that PW and gene expression might also have nonlinear relationships. We fit natural cubic splines with 3 knots and identified 46 DEGs associated with PW in covariate adjusted models (FDR<0.05, **Table S5**, **Figure S8**). The strongest association was with *NCAPH*. We considered the potential role of sex as an effect modifier through sex-stratified analyses and identified 0 DEGs associated with PW in males and 154 DEGs in females (FDR<0.05, **Table S6**), of which the strongest association was with *PSG11*.

We also investigated placental efficiency using a model of BW adjusted for the nonlinear effect of PW (BW_adj_). We identified 458 genes associated with BW_adj_ (**Table S5**; **Figure 2A**). Of these, 7 DEGs were significantly and concordantly associated with BW:PW and BW_adj_ (**Figure 2B**).

Generally, genes were concordantly up- or downregulated in both the BW_adj_ and BW:PW analyses. Thus, the BW:PW ratio and BW_adj_ are correlated metrics of placental efficiency with concordant associations with placental gene expression. We also considered the potential role of sex as an effect modifier using sex-stratified models, and identified 12 DEGs associated with BW_adj_ in males, and 1282 DEGs in females (FDR<0.05, **Table S6**).

Using a “Meet in the Middle” approach, we identified 6 genes associated with both prenatal phthalate exposure (MCINP) and BW_adj_ (**Figure 3A,B**). These genes were all positively associated with MCINP and negatively associated with BW_adj_. We performed a formal mediation analysis for the 6 intersecting genes to investigate if these genes could explain part of the relationship between MCINP exposure and BW_adj_ (**Figure 3C).** Of 6 genes associated with both MCINP and BW_adj_, 4 genes (*DPH1*, *SMO*, *SHROOM2*, *SSBP3*) were identified as significant mediators (*p*<0.05). We also investigated the transcriptome as a high dimensional mediator (**Table 2**), identifying *KIF26B*, *NRP2*, and *FREM1* as significant consistent mediators of the inverse relationships between BW_adj_ and MCIOP, MCMHP, and MCPP, respectively. We did not identify consistent gene-level mediators of the relationships between MCIOP, MCMHP, or MCPP and BW:PW.

**Figure 3:**
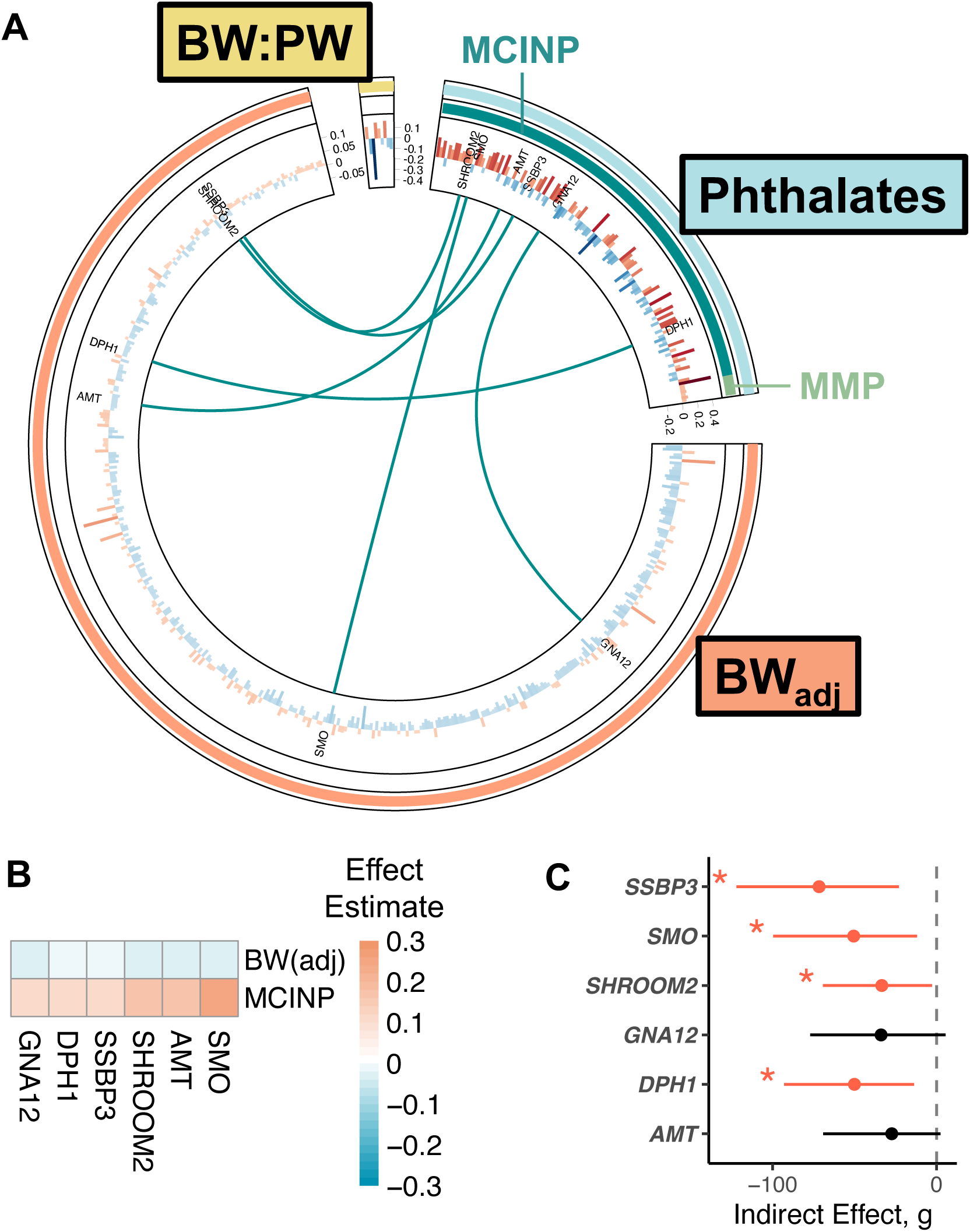
Genes associated with phthalates and/or placental efficiency. Genes were significant at FDR<0.05 for all analyses. A) Circos plot depicts all differentially expressed genes (FDR<0.05) associated with phthalate metabolites, birthweight:placental weight ratio (BW:PW), or birthweight adjusted for placental weight (BW_adj_). Meet in the middle (MITM) genes were associated with both a phthalate and a measure of placental efficiency. MITM genes are labeled and indicated by an arc. B) MITM genes were associated with the phthalate metabolite MCINP and with BW_adj_. C) The point estimates and 95% confidence intervals for the indirect effects of MITM genes in mediating the relationship between MCINP exposure and BW_adj_. *, *p*<0.05.

**Table 2:**
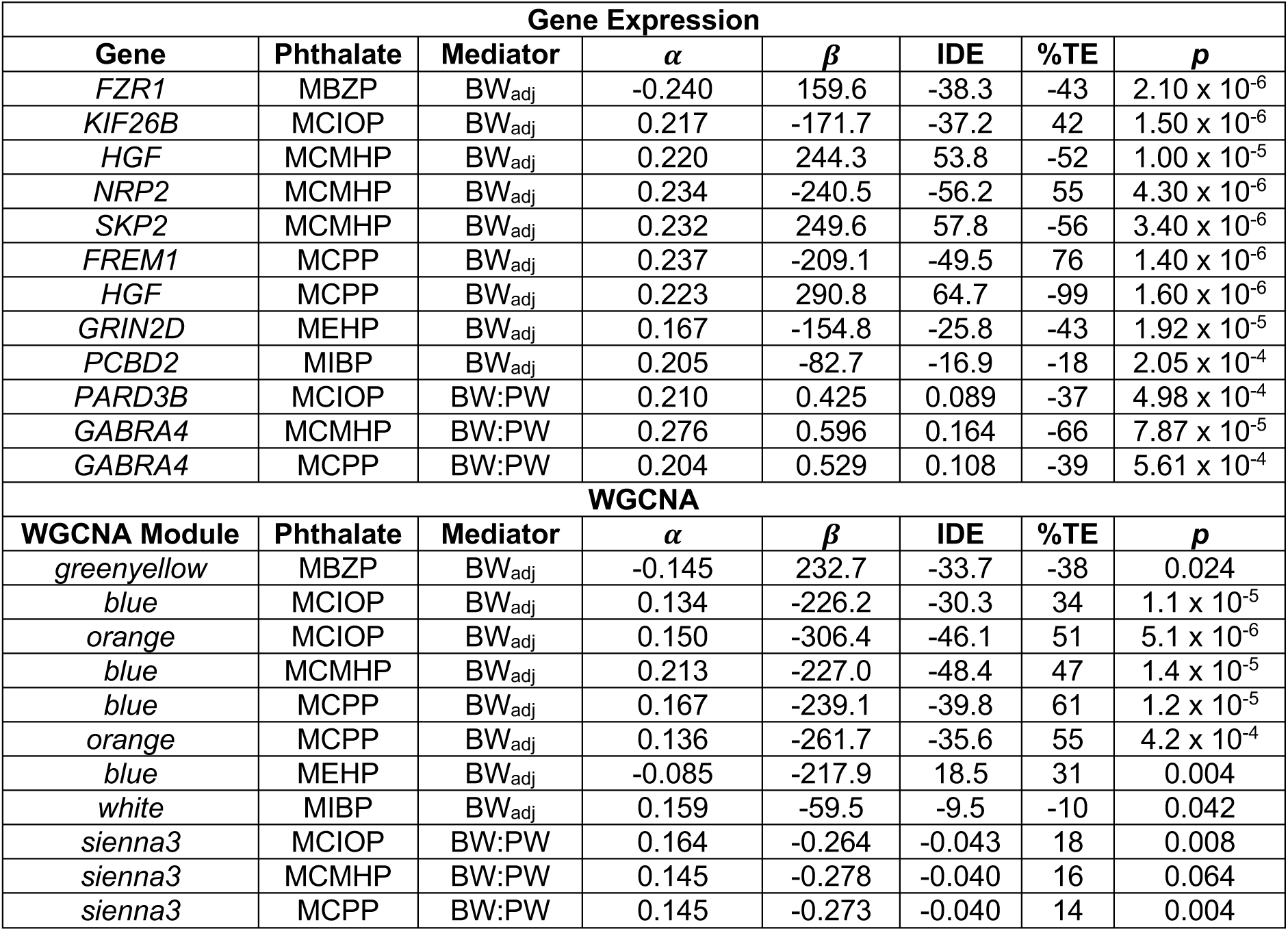
Results of High-Dimensional Mediation for genes and gene modules.

### 4. Network integration of transcriptomic signatures

To investigate the shared and distinct effects of phthalate exposure and placental efficiency on placental gene expression in a network context, we generated co-expressed gene clusters on the full PWG transcriptome dataset (N=465) using Weighted Gene Co-Expression Network Analysis (WGCNA) (*47*). We performed hierarchical clustering based on cluster mean averages and detected 38 gene modules, which represent groups of co-expressed genes. We identified 62 KEGG pathways enriched for genes within 24 of these modules using pathway over-representation analysis (FDR<0.2). We then generated linear regressions to identify WGCNA modules associated with either prenatal phthalate exposure, BW_adj_ or BW:PW using the same covariates as in our gene level analyses (**Figure S5**), and models were considered statistically significant with *p*<0.05, in alignment with previous studies (*48–55*). We identified 27 modules whose expression was associated (p<0.05) with 13 phthalate metabolites (**Figure 4A**). MCINP was associated with the most modules, including 7 positive associations and 9 negative associations. The *green* module was associated with 4 phthalate metabolites, while *turquoise*, *sienna3*, *orange*, *darkgreen*, and *blue* were associated with 3 phthalate metabolites. Overall, 12 modules were associated with multiple phthalate metabolites and 7 of these were directionally concordant. 22 of the phthalate-associated modules were enriched for KEGG pathways (**Figure 4B**). One module (*lightcyan*) was negatively associated (*p*<0.05) with the phthalate metabolite mixture (ß= -0.013). There were 6 metabolites with positive mixture weights and 9 with negative mixture weights, with MEHHP and MEOHP being the most positively and negatively weighted metabolites, respectively (**Table S4**). The *lightcyan* module was enriched for 11 pathways (FDR<0.05) with most related to cell cycle and DNA replication processes (**Figure 4B**).

**Figure 4:**
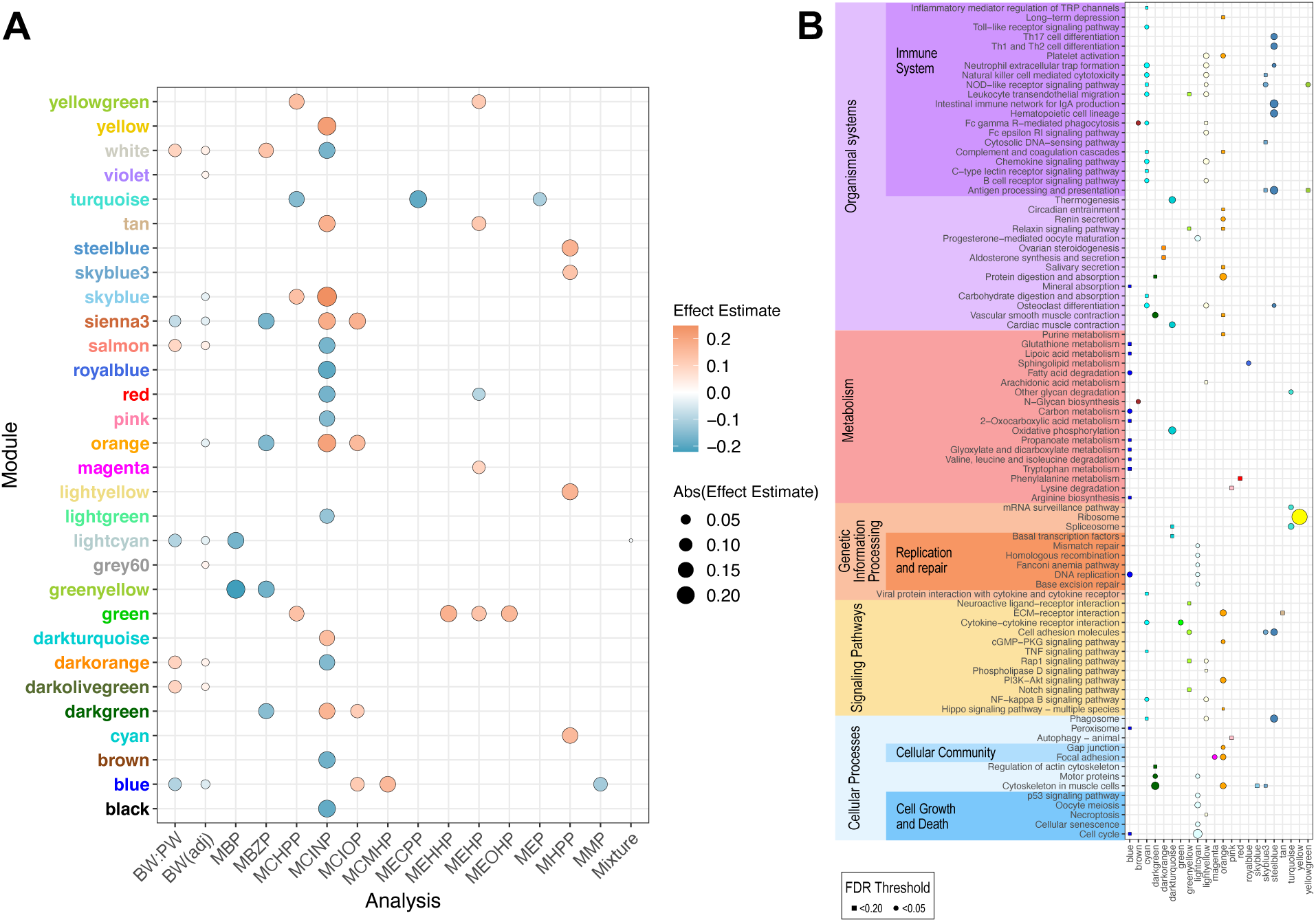
**(A.)** Co-expressed gene modules associated (p<0.05) with birthweight adjusted for placental weight (BW_adj_), BW:PW, and phthalate metabolites. Effect estimates represent a standard-deviation change in eigengene expression for each 100 g increase in birthweight, 1 unit increase in BW:PW ratio or natural log transformed phthalate concentration (ng/mL). The size and color of the dot highlight the strength and direction of the association. **(B.)** Over representation analysis identified KEGG pathways enriched within WGCNA modules. Point size corresponds to -log10(p-value).

We identified 3 modules (*blue*, *lightcyan*, *sienna3*) inversely associated with both BW:PW and BW_adj_ and 4 modules (*darkolivegreen*, *darkorange*, *salmon*, *white*) positively associated both BW:PW and BW_adj_. Additionally, BW_adj_ was significantly inversely associated with two modules (*orange*, *skyblue*) and significantly positively associated with two modules (*grey60*, *violet*). Finally, we identified 4 modules (*blue*, *lightcyan*, *lightgreen*, and *midnightblue*) that were nonlinearly associated with PW (*p*<0.05, **Figure S8B**) using a natural cubic spline.

8 gene modules (*white*, *skyblue*, *sienna3*, *salmon*, *orange*, *lightcyan*, *darkorange*, and *blue*) were associated with BW_adj_ and prenatal urinary concentrations of phthalate metabolites, with all but *orange* and *skyblue* also associated with BW:PW (**Figure 4A**). The directionality of these associations generally varied across multiple phthalate metabolites. The *blue*, *lightcyan*, *orange*, and *sienna3* modules were negatively associated with BW_adj_ and were also negatively associated with phthalate metabolites including MBP, MBZP or MMP. The *white*, *salmon*, and *darkorange* module were positively associated with BW_adj_, and were all negatively associated with MCINP.

Of these shared modules, 5 were enriched for KEGG pathways. The *blue* module was enriched for pathways involved chiefly in metabolism; the *darkorange* module enriched pathways were involved in steroid hormone synthesis and secretion; the *orange* module enriched pathways were involved in cellular adhesion, cytoskeletal function, and extracellular matrix; the *lightcyan* module was enriched for pathways related to cell cycle and DNA replication; and the *skyblue* module was enriched for the Cytoskeleton in Muscle Cells pathway (FDR<0.2, **Figure 4B**). 8 of the overlapping modules (*sienna3*, *blue*, *white*, *lightcyan*, *darkorange*, *salmon, skyblue, orange*) contained genes associated with phthalates, BW_adj_, or BW:PW (**Table S7**).

### Analysis

We conducted high dimensional mediation analysis (HIMA) to evaluate all WGCNA-derived gene modules as mediators (**Table 2**). The *blue*, *greenyellow*, *orange*, and *white* modules were identified as mediators of the relationships between individual phthalates and BW_adj_. In this analysis, we identified consistent mediators, which have total effects and indirect effects with the same sign. We also identified inconsistent mediators, which have total effects and indirect effects with different signs. The *blue* module was identified as a consistent mediator for 4 phthalates, explaining 31-61% of the inverse relationship between MCIOP, MCMHP, MCPP, and MEHP and BW_adj_. All four modules mediating the relationship between phthalates and BW_adj_ contained genes associated with phthalates and BW_adj_ (**Table S7**). The *sienna3* module was a consistent mediator explaining 14% of the inverse relationship between MCPP and BW:PW and 18% of the inverse relationship between MCIOP and BW:PW. The *sienna3* module contained 13 BW_adj_ associated genes and 2 MCINP-associated genes, but no genes in the *sienna3* module were associated with BW:PW (**Table S7**).

## Discussion

This study reveals a potential role of the placental transcriptome as a link between prenatal phthalate exposure and placental efficiency using novel network integration and mediation approaches in a well-phenotyped pregnancy cohort with robust characterization of EDC exposures. We identified associations between metabolites from two different phthalates and the placental transcriptome, with the highest number of associations with MCINP. We also characterized associations between the placental transcriptome and placental efficiency. We identified 7 genes and 2 gene modules that partially explained the relationship between individual phthalates and BW_adj_ with consistent mediated effects. These gene modules were enriched as regulators of key biological pathways, including syncytialization (*orange*) and metabolism (*blue)*. These processes are essential to fetal growth and development because of the placenta’s role in nutrient supply, hormone production, and detoxification. These findings are significant because they reveal toxicological mechanisms by which phthalates may disrupt fetal growth.

We identified associations between the placental transcriptome and maternal urinary concentrations of individual and mixtures of metabolites from different phthalate compounds. MCINP, a secondary metabolite of diisodecyl phthalate (DIDP), was associated with a substantially higher number of genes (126) compared to MMP, which was only associated with 6 genes. In early 2025, the EPA released a new risk evaluation for DIDP which concluded that DIDP, “poses unreasonable risk of injury to human health to female workers of reproductive age” related to 6 industrial uses of the chemical (*56*). The EPA did not find unreasonable risk for the general population, but their evaluation of exposure pathways was not exhaustive. Our study, which is conducted in a generalizable population of pregnant individuals, demonstrates that there may be alterations to placental gene expression patterns following gestational exposure to DIDP and its metabolites. Future research should consider studying mechanisms of DIDP toxicity in relation to placental biology. Only two genes provisionally associated (p<0.005) with the phthalate mixture (*CEP63*, *RGS4*) were also associated with individual phthalate metabolites, indicating that phthalate mixture exhibits different effects than the compounds individually. Overall, phthalate metabolites have half-lives of less than 24 hours (*3*, *57*), so the urine samples used in this analysis represent a short term exposure window.

We have previously assessed associations between phthalate metabolites and the placental transcriptome in the Conditions Affecting Neurocognitive Development and Learning in Early childhood (CANDLE) study which identified associations between 32 protein coding genes and 4 phthalate metabolites in the full sample (N=760) (*42*). In contrast to this study in PWG participants, the analysis in CANDLE participants separated phthalate measurements by trimester of sample collection, while herein our analysis used a geometric mean across study visits to calculate a single phthalate metabolite concentration summary for each participant, which reduces exposure misclassification (*58–60*). The results of the present study identified substantially more phthalate-associated genes (132 genes), and the two metabolites (MCINP and MMP) with gene associations in PWG did not have any gene associations in CANDLE. The differences in findings across these two studies could be attributed to baseline differences in the location and demographics of the study population with PWG recruiting from Seattle and Yakima, WA, whereas CANDLE was recruited from Shelby County, TN. We adjusted for similar covariates in our linear models to account for sample demographics. Gene expression differences may also be attributable to exposure level differences. Phthalate measurements vary across time of enrollment in these cohorts but are relatively comparable across these cohorts and in alignment with the US-based National Health and Nutrition Examination Survey (NHANES), based on year of enrollment (*61*). Lastly, as previously noted, the substantial differences in quantity of gene associations could be related to the use of a single composite exposure measure (calculated as a geometric mean) for each phthalate in the present study which reduces exposure misclassification compared to the single measures that may have biased the CANDLE results toward the null.

While the placenta is often framed as a simple conduit of fetal nutrients, it is a highly metabolic organ, and we report nonlinear relationships between PW and birthweight and PW and placental gene expression. To holistically capture placental efficiency, we evaluated both BW:PW ratio, which captures the relationship between the interrelated fetal and placental growth trajectories (*62*), as well as birthweight adjusted for placental weight (BW_adj_) a measure of placental efficiency that accounts for placental weight. We have elected not to adjust for gestational age in our models because poor fetal growth and placental insufficiency are risk factors for preterm birth and adjustment may introduce collider bias (*63–65*). We observed that genes and co-expressed gene modules were generally concordantly associated with both metrics of placental efficiency, and we identified 7 concordant DEGs, including genes involved in cytoskeletal function and motility (*KIF26B*, *CAMSAP3*), neuronal development (*NDRG2*, *PRX*), cell cycle (*TOP2A*), and protein degradation (*ASB2*). We also identified 7 concordant co-expressed gene modules, including the *sienna3*, *blue* and *lightcyan* modules which were inversely associated with BW:PW and BW_adj_. While correlated, these measures of placental efficiency captured different variation in gene expression, as BW_adj_ was uniquely associated with 451 genes and 4 modules.

We identified 4 genes (*SMO*, *SHROOM2*, *DPH1*, and *SSBP3*) that mediated the relationship between MCINP and BW_adj_. *SMO* codes for a G protein-coupled receptor that is involved in the activation of the hedgehog signaling pathway, which is critical for morphogenesis during development (*66*). Hedgehog signaling is important for placental development where it has been implicated in epithelial-mesenchymal transition which is essential for the formation of the syncytiotrophoblast (*67*). Though there are no known associations between *SMO* and birthweight outcomes, *SMO* expression is increased in LNCaP cells exposed to MEHP, but DNA methylation is not altered (*68*, *69*). *SSBP3* encodes a transcriptional coactivator that regulates head development and fetal body growth in mice (*70*), and regulates trophoblast-like cell differentiation in mouse embryonic stem cells (*71*). There is limited information about the association of *SHROOM2* and *DPH1* with placental development, birthweight, or phthalate exposure. However, all four genes were differentially expressed in HTR-8/SVneo placental cells exposed to MEHP with *DPH1* showing increased expression and *SMO*, *SHROOM2* and *SSBP3* showing decreased expression, which is opposite in directionality to the response to MCINP that we saw in the present study (*72*). To date, most toxicological studies of phthalates in the placenta have been conducted with MEHP or its parent compound, DEHP (*20*, *73*), so more research is needed to evaluate the toxicological response to DIDP metabolites, like MCINP, which may be different from MEHP.

Using a HIMA approach, we also identified a handful of gene-level mediators linking phthalate exposure to BW_adj_. These included genes encoding an intracellular motor protein (*KIF26B*), a VEGF receptor (*NRP2*), and a basement membrane adhesion protein (*FREM1*) as mediators of the relationship between BW_adj_ and MCIOP, MCMHP, and MCPP, respectively. With HIMA, we identified the *blue* and *orange* modules as consistent mediators with the relationship between BW_adj_ and MCIOP, MCMHP, MCPP, or MEHP, which mirror the gene-level mediation findings. Paired with omics data, mediation analysis is a powerful analytical framework to investigate biological mechanisms. However, mediation analysis has strong assumptions, including no unmeasured confounding. Our mediation analysis should be interpreted as hypothesis generating to guide further research to better understand the relationship between prenatal phthalate exposure and placental efficiency. We also employed network methods to better understand the biological pathways that link prenatal phthalate exposure to BW_adj_ and BW:PW, and identified several converging findings.

Several WGCNA modules (*orange*, *blue, lightcyan*, *darkorange* and *sienna3*) emerged as consistent mediators between phthalate metabolites and measures of placental efficiency, and genes within these modules were involved in key biological pathways. The *orange* module was enriched for genes in cytoskeleton, cellular adhesion, and extracellular matrix pathways. We have also identified these pathways as associated with phthalate exposure in other contexts, including in the CANDLE cohort, where MCIOP and MMP were associated with gene expression changes in the adherens junction pathway (*42*), and *in vitro*, where MEHP treatment upregulates cytoskeletal regulation in primary placental trophoblasts and HTR-8/SVneo cell lines (*72*). These shared changes might be related to the process of syncytialization, when components crucial for cytotrophoblast fusion are activated (*74*), the extracellular matrix is altered (*75*), and cytotrophoblasts adhere to the existing syncytial layer to form a large, multinucleated barrier (*76*). This process is important to placental function and fetal growth because the syncytiotrophoblast plays a crucial role in maintaining gas and nutrient exchange (*74*, *77*), and produces key hormones such as chorionic gonadotropin hormone (hCG) (*78*, *79*), and impaired markers of syncytialization have been linked to intra-uterine growth restriction (*80*). *In vitro* studies have revealed that phthalates can disrupt trophoblast differentiation (*18*), and inhibit trophoblast invasion (*19*), but the effects on the process of syncytialization have not been reported. Together, this work suggests that phthalates may impair syncytialization leading to altered placental growth, with long term implications to fetal development and growth.

The *blue* module was also a consistent mediator between prenatal exposure to MCIOP, MCMHP, MCPP, and MEHP and BW_adj_, and was enriched for genes in 10 different metabolic pathways, including metabolism of different amino acids as well as one carbon metabolism. Metabolism is relevant to placental growth as the placenta is a highly metabolic organ that needs tremendous energy and resources to support its own growth as well as the fetus in a 9 month timespan (*81*). Changes in the placental metabolome have been linked to fetal growth restriction, including decreased concentration of amino acids (*82*), other metabolites (*83*), and recent work has suggested that alterations in one carbon metabolism may affect fetal growth through perturbed DNA methylation (*84*). Prenatal phthalate exposure has been associated with differences in the fetal, placental, and maternal metabolome (*85–88*). Intriguingly, one recent study used a MITM approach to reveal a subset of metabolites (including tryptophan) that were associated with both prenatal phthalate exposure and newborn neurobehavior. Other work by this group using a MITM approach on neonatal blood identified revealed shared metabolic signatures related to prenatal exposure to other environmental chemicals (per- and polyfluoroalkyl substances) and fetal growth (*89*) as well as gestational length (*90*). More work is needed to investigate how phthalates disrupt these metabolic pathways and the contribution of these pathways to fetal growth.

At low and high PW, the placenta shows higher expression of the *lightcyan* gene module, which was also associated with BW_adj_. While placental expression of genes within the *lightcyan* module were not associated with any individual phthalate, it was negatively associated with the phthalate mixture. The *lightcyan* module was enriched for cell growth pathways (cell cycle, cellular senescence), as well as replication and repair pathways (DNA replication, base excision and mismatch repair, and homologous recombination). Together, this may suggest that phthalates induce DNA damage resulting in cellular senescence. Phthalates can induce dsDNA breaks and inhibit the DNA repair pathways needed to respond to these breaks *in vitro* (*21*, *91–93*). DNA damage is linked to a number of pregnancy complications (*94*), including elevated birthweight (*95*) and inadequate fetal growth. Based on this body of work, we hypothesize that DNA damage may contribute to impaired cell growth, which may also be involved in the process of syncytialization (*96*). Together, this work highlights converging mechanisms related to both phthalate exposure and impaired placental efficiency.

Our WGCNA analysis validates known mechanisms of phthalate toxicity, while also revealing novel mechanisms and genes that may not be members of established pathways. The *darkorange* module was positively associated with BW:PW and BW_adj_ and was negatively associated with MCINP. This module was enriched for genes involved in steroid synthesis, which is a well-established mechanism of phthalate toxicity (*97*, *98*). These results replicate prior findings in CANDLE, in which we observed positive associations between prenatal MMP exposure and the steroid biosynthesis pathway. The *sienna3* module emerged as an important mediator and our MITM analysis revealed negative associations between this gene module and BW:PW and BW_adj_ along with associations with multiple phthalate metabolites. With 43 genes, the *sienna3* module was not significantly enriched for any pathways after FDR correction. However, *sienna3* contained 13 DEGs inversely associated with BW_adj_, including genes involved in cell adhesion and extracellular matrix (*CADM3*, *COL14A1*, *HAPLN1*, *LAMA2*, *PCDH10*, *PCDH18*, *SPON1*). This module also contained *CXCL14*, which inhibits trophoblast invasion and high umbilical cord expression has been associated with low birthweight (*99*, *100*). Overall, genes and pathways involved in syncytialization, metabolism, DNA damage and cellular senescence, and steroid biosynthesis were repeatedly identified across our MITM and HIMA approaches to mediation. Biological pathways and cellular functions identified in both these approaches support biological importance and warrant further study *in vitro* and independent cohorts.

This analysis should be interpreted in light of inherent limitations with this study. Gene expression was quantified using bulk RNA sequencing data and thus does not account for differences in the distribution of cell types collected within each sample, which is a well-established challenge of all bulk RNA sequencing analyses of the placenta (1, 2). However, this study makes an important contribution to this field as the first study to mechanistically link prenatal phthalate exposure and placental efficiency using high dimensional mediation and network analyses. Strengths of this study include that we considered mixtures of phthalate metabolites, which are a more robust characterization of real-world exposures. We have also deployed high dimensional mediation analysis (5–7), which is an advancement beyond meet in the middle analysis-based approaches that have been used to integrate omics data in the context of other exposure-outcomes studies (8–10). To our knowledge, this is the first study to apply these approaches to placental efficiency, which is a highly relevant outcome to EDC exposures and is directly related to the placental transcriptomics data generated in this study.

In summary, this study reveals potential mechanisms linking the relationship between prenatal phthalate exposure and placental efficiency through placental gene expression. Our results highlight specific mechanisms including syncytialization, metabolism, DNA damage and cellular senescence, and steroid biosynthesis. Further experimental work as well as external validation is needed to confirm these findings. Placental efficiency is an important regulator of fetal development and has implications for infant and childhood growth outcomes, which warrants further study.

## Materials and Methods

### 1. Study participants

Placental samples were collected from pregnant women in Seattle and Yakima WA as part of the Global Alliance to Prevent Prematurity and Stillbirth (GAPPS) study. Over 600 of these pregnant women and their children subsequently enrolled into the nationwide NIH ECHO study as the PWG consortium in 2016 (**Figure S1**), with placental transcriptomics data available on 465 participants. Inclusion criteria for the GAPPS study included: participant age of 18 years of age or older or medically emancipated and confirmed to be pregnant by self-test or by physician’s medical testing. In this analysis, we included all participants with RNA sequencing data to generate the WGCNA modules. For further analyses, we excluded participants with placental abruption (*N*=14) or multi-fetal gestations (*N*=13) in this analysis (**Figure S1**). For phthalate analyses, our final sample included *N*=222 participants with prenatal urinary phthalate measurements and complete covariate data. For placental efficiency analyses, our final sample included *N*=253 participants with birthweight, placental weight, and covariates. Mediation analyses included *N*=163 participants included in both phthalate and placental efficiency analyses (**Figure S1**). The Seattle Children’s Research Institute IRB approved all research activities for the GAPPS cohort. (IRB # STUDY00000608)

### 2. Collection of maternal urine and quantification of phthalate metabolite concentrations

Maternal urine was collected using phthalate-free polypropylene containers at one of two clinical visits and stored at −80°C. Samples were analyzed for 21 phthalate metabolites using solid-phase extraction and high-performance liquid chromatography tandem mass spectrometry as previously described (*6*, *101–103*). Field, process, and instrument blanks were included for quality control. Specific gravity was determined using a handheld refractometer to account for urinary dilution (*104*). Only those phthalate metabolites that were detected in >70% of samples were included in the analysis (**Table S2**). DEHP was calculated as the molar sum of 5 of its monoester metabolites as follows: DEHP = (MEHP * (1/278.34)) + (MEHHP * (1/294.34)) + (MEOHP * (1/292.33)) + (MECPP *(1/308.33)) + (MCMHP*(1/308.33)). For samples below the limit of detection (LOD), the concentration was reported as the LOD/√2. Metabolites were natural log transformed to obtain normal distributions. For samples with measurements at multiple visits, we averaged the average values across multiple timepoints using a geometric mean as in previous studies (*105–107*). Separate phthalate datasets for 2^nd^ trimester (*N*=139) and 3^rd^ trimester (*N*=189) samples were also prepared for subsequent analyses. A summary of visit timing and samples per participant is available in **Figure S2A,B**.

### 3. Placental sample collection

Placental weight and birthweight were measured by the GAPPS placental biorepository at the time of sample collection from the medical record. Prior to weighing, placentas were trimmed to remove fetal membranes and the umbilical cord was cut to approximately 5 cm from the insertion site. Four separate vertical placental tissue punches were taken at 2 sites 7 cm apart at a depth of 8 mm, and stored in RNAlater within 30 minutes of delivery and stored at -20 °C, as previously described (*108*). Specimens then were shipped to the GAPPS facility and stored at -80 °C. Fetal villous tissue was manually dissected and cleared of maternal decidua using standard protocols developed by the GAPPS placental biorepository. The BW:PW ratio, an indicator of placental efficiency, was calculated as the ratio of birthweight (g) to trimmed placental weight (g).

### 4. Placental sample processing and RNA sequencing

Placental tissue processing, RNA isolation and sequencing have previously been described (*108–110*). Briefly, 30 mg of fetal villous placental tissue was homogenized in 600 µl buffer RLT Plus with β-mercaptoethanol using a TissueLyser LT instrument (Qiagen; Germantown, MD). RNA was isolated using the AllPrep DNA/RNA/miRNA Universal Kit (Qiagen, Germantown, MD) according to the manufacturer’s recommended protocol. RNA integrity was determined with a Bioanalyzer 2100 using RNA 6000 Nanochips (Agilent; Santa Clara, CA), and only RNA samples with an RNA Integrity Number (RIN) >7 were sequenced. Samples were sequenced to an approximate depth of 30 million reads by either Illumina HiSeq 4000 or Novoseq at the University of Washington Northwest Genomics Center. To ensure replicability between these platforms, 28 samples were run across both platforms, and the results were highly correlated (correlation coefficient 0.9, p=4.48x10^-11^). RNA sequencing quality control was performed using both the FASTX-toolkit (v0.0.13) and FastQC (v0.11.2) (*111*). Transcript abundances were estimated by aligning to the GRCh38 transcriptome (Gencode v33) using Kallisto (*112*), then collapsed to the gene level using the Bioconductor tximport package, scaling to the average transcript length (*113*). Only protein-coding genes were included in this analysis. Genes were filtered to remove low-expressing genes with average logcpm<0.

### 5. Identification of differentially expressed genes (DEGs)

We identified differentially expressed genes and gene modules in models adjusted for selected potential confounders which were identified a priori by reviewing research literature and construction of directed acyclic diagrams, which are presented in **Figure S5**. In our analyses of associations between phthalates and the placental transcriptome, we adjusted for urine specific gravity (geometric mean), RNA sequencing batch, fetal sex, maternal BMI, maternal race, maternal ethnicity, maternal education, maternal age, maternal smoking (via self-report or urinary cotinine detection), parity, delivery method, and labor type. In our analyses of associations between the placental transcriptome and BW:PW, we adjusted for the same confounders in addition to gestational diabetes and gestational hypertensive disorders (including either preeclampsia, gestational hypertension, or chronic hypertension). In our analysis of birthweight adjusted for placental weight (BW_adj_) as a measure of placental efficiency, we adjusted for the same confounders as in the BW:PW model and included a nonlinear term for placental weight. We conducted a sensitivity analysis without the two participants with the highest placental weights to determine if they were driving the nonlinear association between PW and BW and found the results were consistent with a nonlinear relationship (**Figure S3C**). We additionally evaluated *P*-value distributions from additional sensitivity analyses for the nonlinear relationship to determine if the two samples with the highest placenta weights were driving the nonlinear association with gene expression or if linear modeling was appropriate (**Figure S9**). These results supported evaluating a nonlinear relationship between PW and gene expression. Preterm newborns are more likely to be growth restricted relative to fetuses remaining in utero (*63*, *114*), such that gestational length might act as a collider in the analysis of placental efficiency and placental gene expression. Gestational length was excluded as a mediator on the causal pathway between phthalate exposure and both BW_adj_ and BW:PW (*115*). We adjust for race and ethnicity through the framework that these variables are social constructs reflecting decreased opportunities, increased experiences of racism, and exposure to environmental pollutants for minorities and women of color (*116*, *117*). To assess effect modification by child sex, we performed a sex-stratified analysis and adjusted for all variables described above.

Differentially expressed genes associated with the phthalates, PW, BW:PW, or BW_adj_ were identified using the limma-voom pipeline, which incorporates observation-level weights to account for relationship between the mean and variance of the log CPM values (*118*). To capture the nonlinear relationship between PW and gene expression, we used natural cubic splines (*splines::ns* function) with 3 knots placed at the 25^th^, 50^th^, and 75^th^ percentiles for PW. We performed adjustment for multiple comparisons using the Benjamini-Hochberg approach (*119*), and genes were considered statistically significant at a false discovery rate (FDR) <0.05. We examined the association between the mixtures of phthalate metabolites and the placental transcriptome at a gene and gene module level using quantile g-Computation (QGComp) (*120*), adjusting for the same covariates as above, except for smoking which did not have enough contrast to be included in the mixture models. We used the ewas_qgcomp function which was recently developed for high dimensional outcomes including RNA sequencing and DNA methylation data (*121*). Bootstraps (*N*=1000) were used to estimate confidence intervals for the sum effect estimate associated with a one quartile higher concentration of every component of the metabolite mixture. DEHP was excluded from the mixtures analysis which included only phthalate metabolites. For our mixtures analysis, individual genes were considered provisionally significant at p<0.005 (*122*). For all phthalate results, effect estimates in tables and figures represent the beta coefficient from respective models.

### 6. Integrating signatures using weighted gene co-expression network analysis

Weighted gene co-expression network analysis (WGCNA) was performed on the full RNA-sequencing data set (*N* = 465). Filtered count data was normalized using conditional quantile normalization (cqn::cqn function) to gene length and CG composition (*123*). WGCNA was conducted using the WGCNA package (version 1.72-1) (*47*) as an unsigned network constructed using Pearson’s correlation, hierarchical clustering based on cluster mean averages, and modules containing at least 20 genes. Modules were determined using dynamic tree cut (WGCNA::cutreeDynamic). Genes assigned to a given module that were highly correlated with that module’s eigengene (|*r*|>0.8) were selected as the given module’s hubgenes (**Table S8**). Modules were characterized by conducting KEGG pathway (release 111.0) over-representation analysis (limma::kegga function) on gene members, and pathways were considered significantly enriched when FDR<0.2 (**Table S9**)

Multiple linear regression was used to identify WGCNA modules associated with phthalate metabolites, PW, BW_adj_, or BW:PW. These models were adjusted for the same covariates as in the gene-level analysis using natural cubic splines for PW as described and were considered statistically significant with p<0.05, in alignment with previous studies (*48–55*). We examined genes and gene modules that were associated with both phthalate metabolites, BW_adj_, and BW:PW using a “meet in the middle analysis”, which refers to finding the intersection between markers of exposure and markers of outcome (*124*). We then performed a series of mediation analyses on the overlapping DEGs. In these mediation analyses, the dependent variables were BW:PW or BW_adj_, the mediation variable was the expression (logCPM) of the candidate mediator gene or the eigengene value, and the exposure variable was the maternal urinary phthalate metabolite concentration. All mediation analyses were run using the R “mediation” package. For each candidate mediator gene, we ran 2 models: the fitted model between exposure (phthalate metabolite) to mediator (gene expression of candidate gene), and then the fitted model between the mediator (gene expression of candidate gene) and outcome (BW:PW or BW_adj_). Causal mediation effects were estimated using the mediate function, with non-parametric bootstrapping performed across 10,000 simulations. All models were adjusted for the same variables as in the initial analysis, including natural cubic splines for PW fit as described.

### 7. High dimensional mediation analysis

As a high-dimensional mediator, genes within the transcriptome are highly correlated, which can introduce bias. We employed the high-dimensional mediation analysis (HIMA) approach implemented in the R package *hima* (*125*) using gene expression data from the full transcriptome and WGCNA eigengenes. In HIMA, potential mediators are selected through sure independent screening, variable selection is conducted with minimax concave penalty, and joint significance testing is conducted for mediation effects. Models were adjusted for covariates and FDR was set at the 0.05 level.

### 8. Statistical methods

Linear regression models were used to evaluate the association between phthalates and BW_adj_ and BW:PW. Models were adjusted for confounding and precision variables (**Figure S4E**) and included 163 participants. Associations were considered significant at *p*<0.05.

## Acknowledgments

This research was conducted using specimens and data collected and stored on behalf of the Global Alliance to Prevent Prematurity and Stillbirth (GAPPS) Repository. We would like to thank the GAPPS participants as well as the research staff and clinicians that facilitated these studies. The content is solely the responsibility of the authors and does not necessarily represent the official views of the National Institutes of Health. This manuscript has been reviewed by PATHWAYS for scientific content and consistency of data interpretation with previous PATHWAYS publications.

## Funding

National Institute of Environmental Health Sciences R01ES033785 (AGP)

National Institute of Health 5UG3OD035508 (SS)

National Institute of Health UH3OD023271 (SS, KL, NB, QZ)

National Institute of Environmental Health Sciences P30ES007033 (JK)

## Author contributions

Conceptualization: MP, SL, SS, AP

Data Curation: MH, DD, KK, JM, TB, AP

Formal Analysis: MP, SL

Funding Acquisition: NB, KL, SS, AP

Investigation: MP, SL, SS, AP

Methodology: JM, TB, MP, SL, AP, AS, KK

Project Administration: CT, MH, SS, AP

Resources: JM, TB, CT, SS, AP

Software: MP, SL, JM

Supervision: SS, AP Visualization: MP, SL, AP

Writing—original draft: MP, SL, AP

Writing—review & editing: MP, SL, JM, TB, MH, DD, KK, CT, NB, KL, SS, AP, AS, QZ

## Competing interests

Authors declare that they have no competing interests.

## Data and materials availability

Placental transcriptomic data for PWG participants is available on DBGAP (phs003620.v1.p1). De-identified covariate data may be available on request with review of a detailed data analysis plan by the PWG Scientific Committee. Access to data will also require approval by the recipient institution’s IRB and a formal data use agreement. Contact the corresponding author for more information. All code is available on a centralized GitHub repository: https://github.com/Paquette-Lab/PWG_PlacentalTranscriptomeBWPWEDCs

## Notes

### Competing Interest Statement

The authors have declared no competing interest.

