## Supplementary Materials for "The Placental Transcriptome Serves as a Mechanistic Link between Prenatal Phthalate Exposure and Placental Efficiency"

Mariana Parenti *et al.*

**This PDF file includes:**

Figs. S1 to S9
Tables S1 to S2

**Other Supplementary Materials for this manuscript include the following:**

Tables S3 to S9 (Excel Files)

**SUPPLEMENTARY FIGURES**

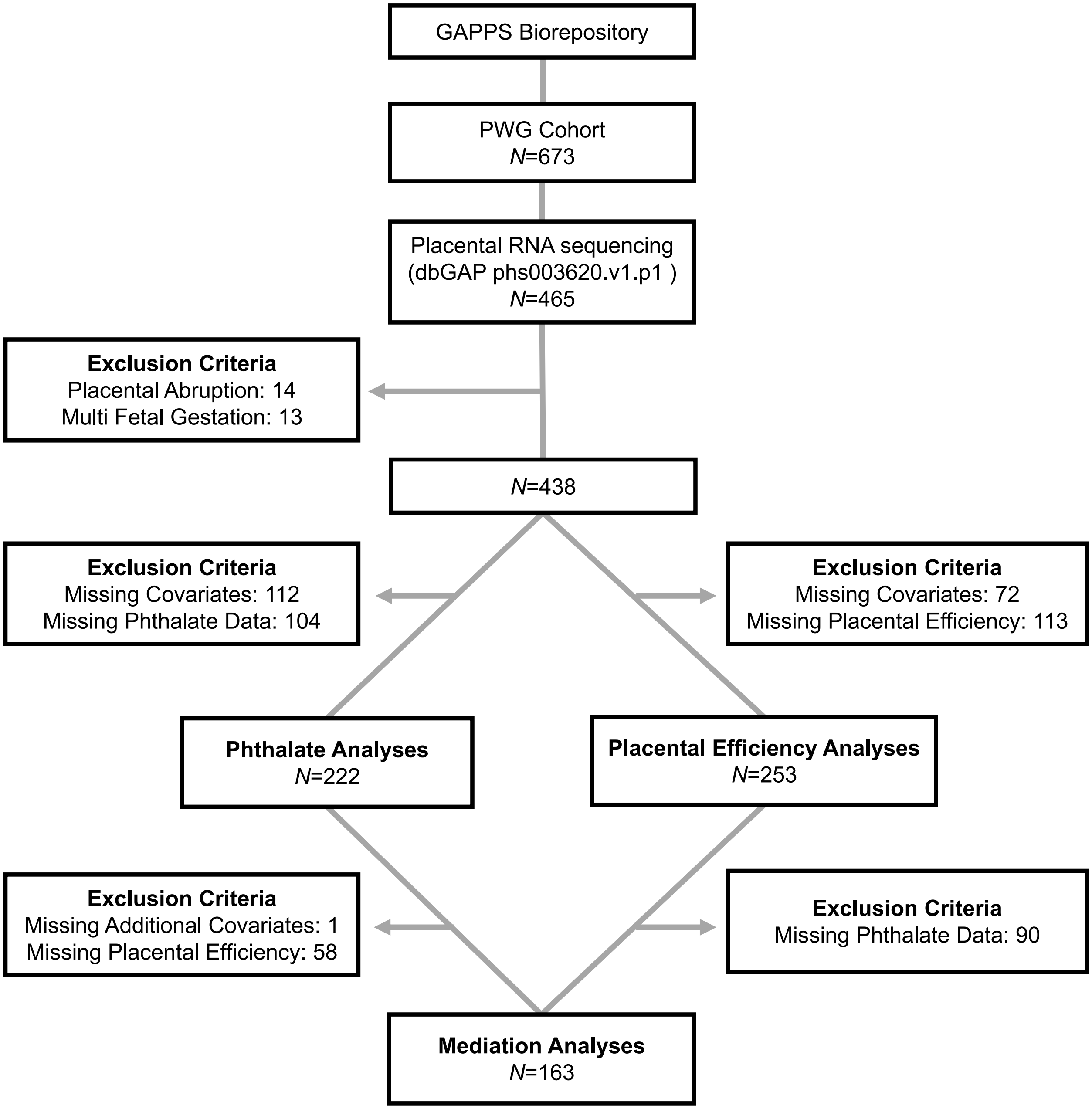


***Figure S1:*** Flowchart of the number of participants included in each analysis based on data availability and inclusion/exclusion criteria.

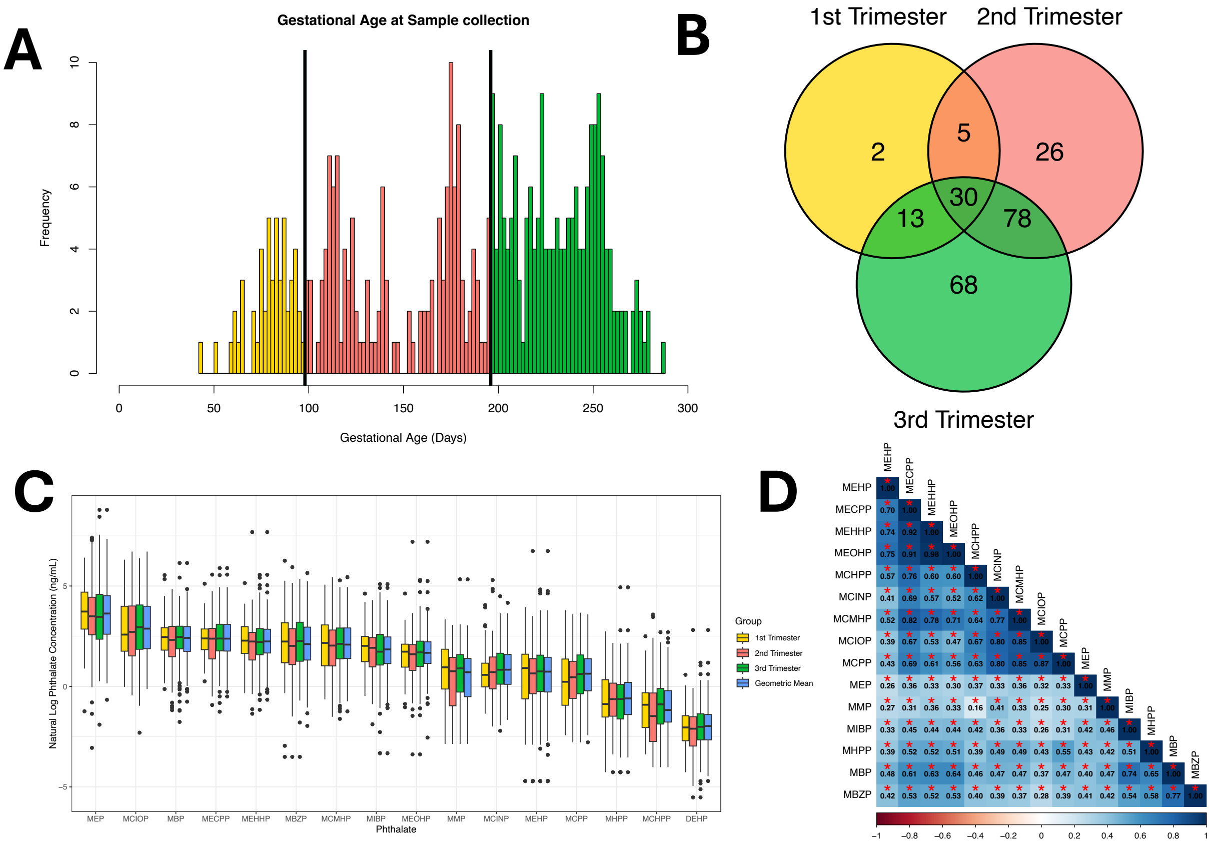


***Figure S2:*** **(A.)** Histogram of gestational age in days for urinary phthalate measurements. **(B.)** Number of participants with phthalate measurements for each trimester. **(C.)** Boxplot of natural log transformed phthalate metabolite concentrations (ng/mL) (N=222). DEHP is expressed as the molar sum of its metabolites and is expressed as natural log transformed µM **(D.)** Pearson correlations between phthalate metabolite natural log transformed concentrations. Significant correlations are designated with an asterisk

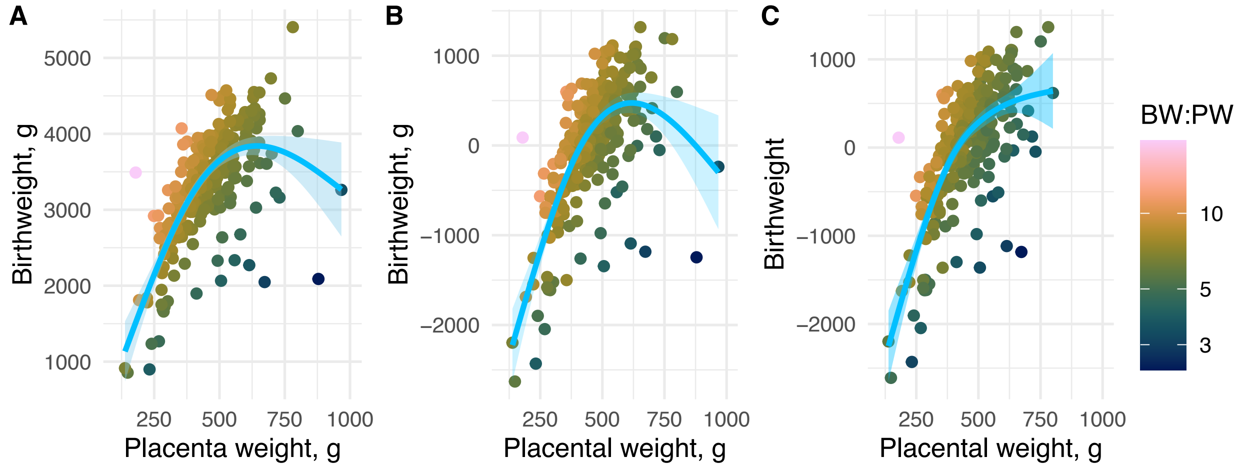


***Figure S3:*** Relationship between birthweight (BW), placental weight (PW), and birthweight:placental weight ratio (BW:PW). A natural cubic spline was fit with knots at the 25, 50, and 75 percentiles. (**A**) In the unadjusted model ($N$=253), the relationship was significant and nonlinear (effective degrees of freedom, EDF=2.86, $p < 2\times 10^{-16}$ ). (**B**) Birthweight is presented as partial residuals adjusted for fetal sex, maternal age, pre-pregnancy BMI, maternal race, ethnicity, education, smoking, parity, gestational diabetes, and hypertensive disorders of pregnancy ($N$=253). The relationship was significant and nonlinear (EDF = 2.88, $p < 2\times 10^{-16}$). (**C**) In a sensitivity analysis with the participants with the two highest placental weights removed ($N$=251), the relationship remained significant and nonlinear (EDF = 2.80, $p < 2\times 10^{-16}$). Birthweight is presented as partial residuals as in B.

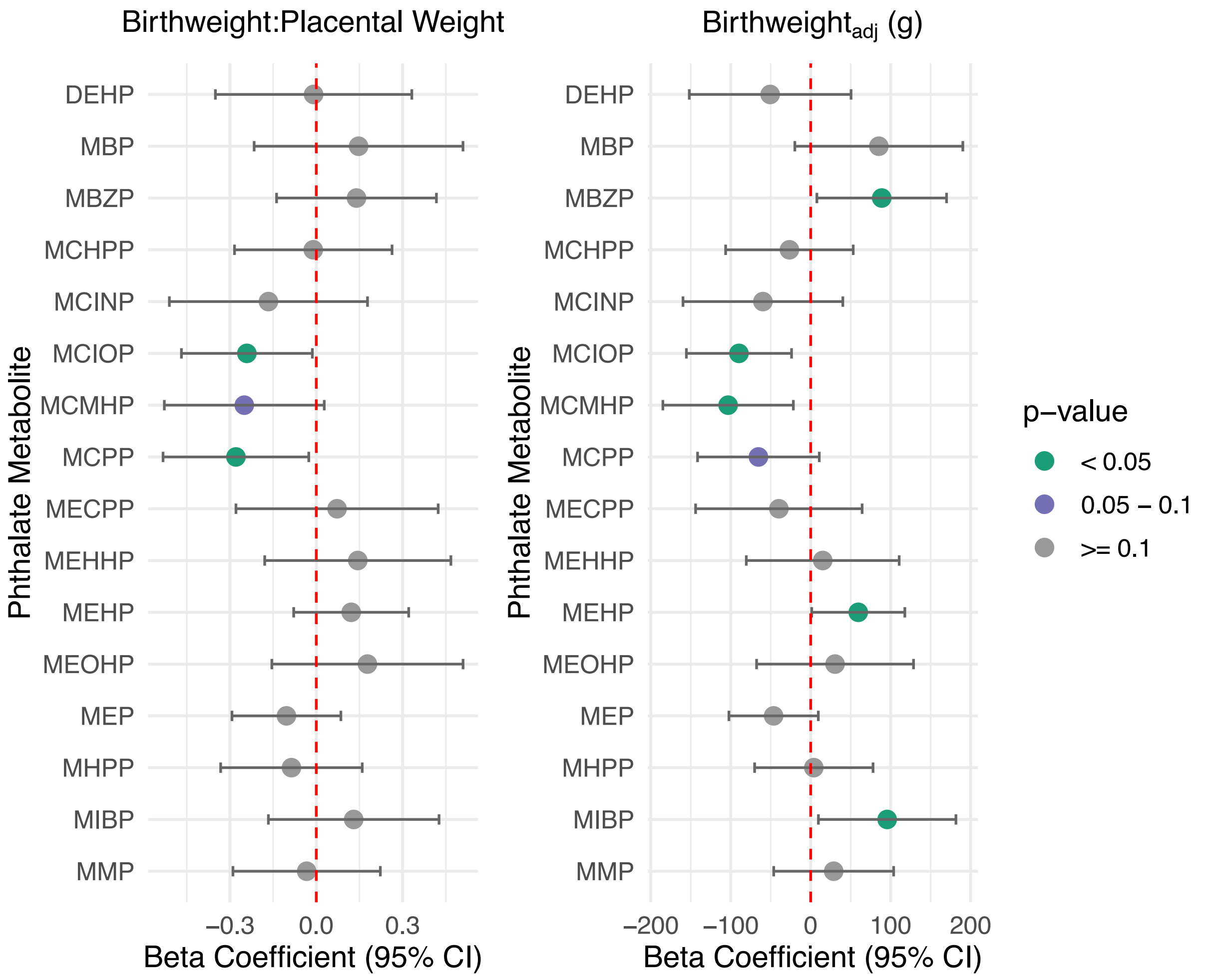


***Figure S4:*** Association between phthalate metabolites, $BW_{adj}$, and BW:PW adjusted for urine specific gravity, maternal age, maternal BMI, maternal race, maternal ethnicity, maternal education, labor type, delivery method, parity, fetal sex, maternal smoking, and placental weight (for $BW_{adj}$ only). Beta coefficients represent the change in BW:PW or $BW_{adj}$ following a one unit increase in natural log transformed phthalate concentrations. Green points indicate significance ($p<0.05$), while purple points indicate marginal significance ($p<0.1$).

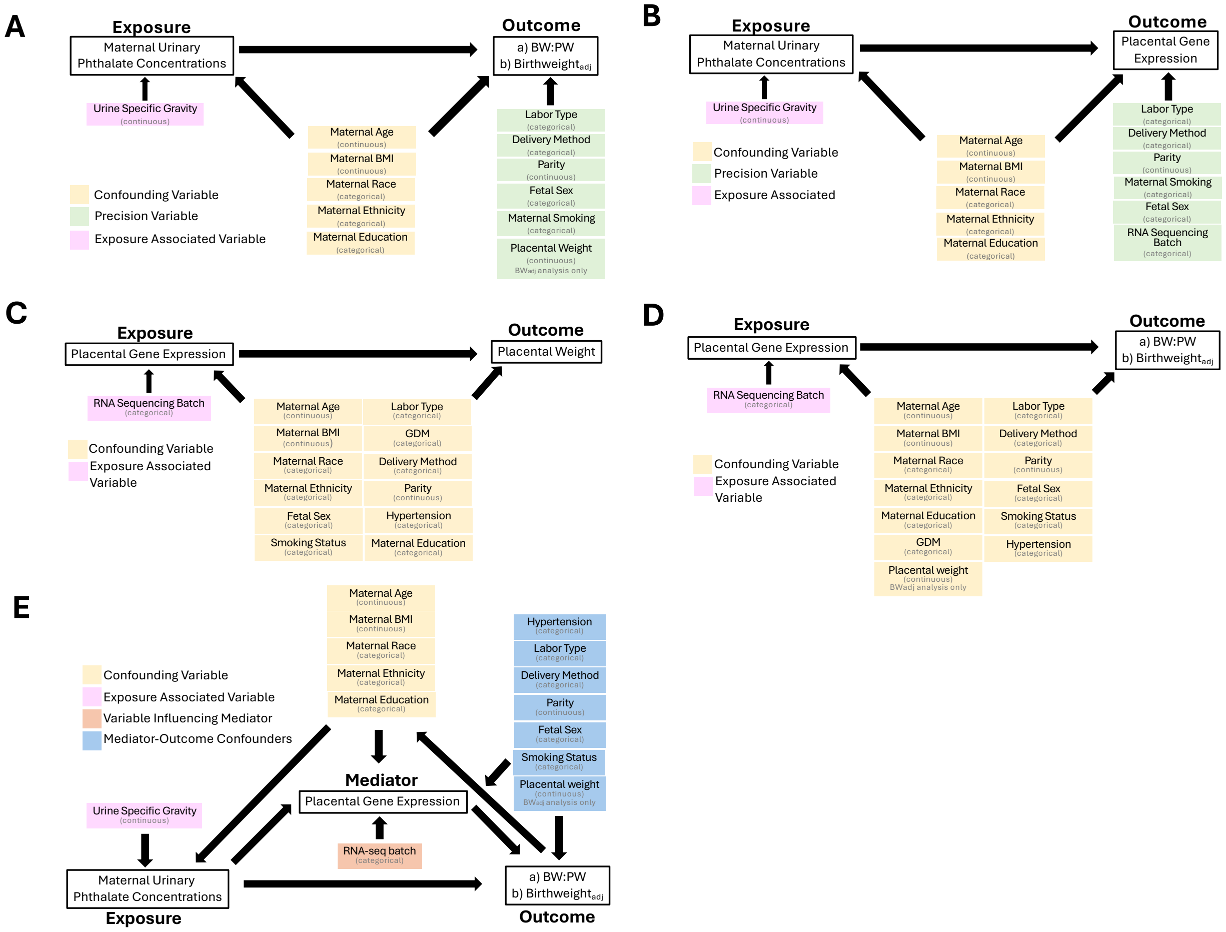


***Figure S5:*** Directed Acyclic diagrams describing adjustment variables for (A) Phthalates and Placental Efficiency, (B) Phthalates and Placental Gene Expression; (C) Placental Weight, (D) Placental Efficiency and Placental Gene Expression, and (E) Mediation Analyses.

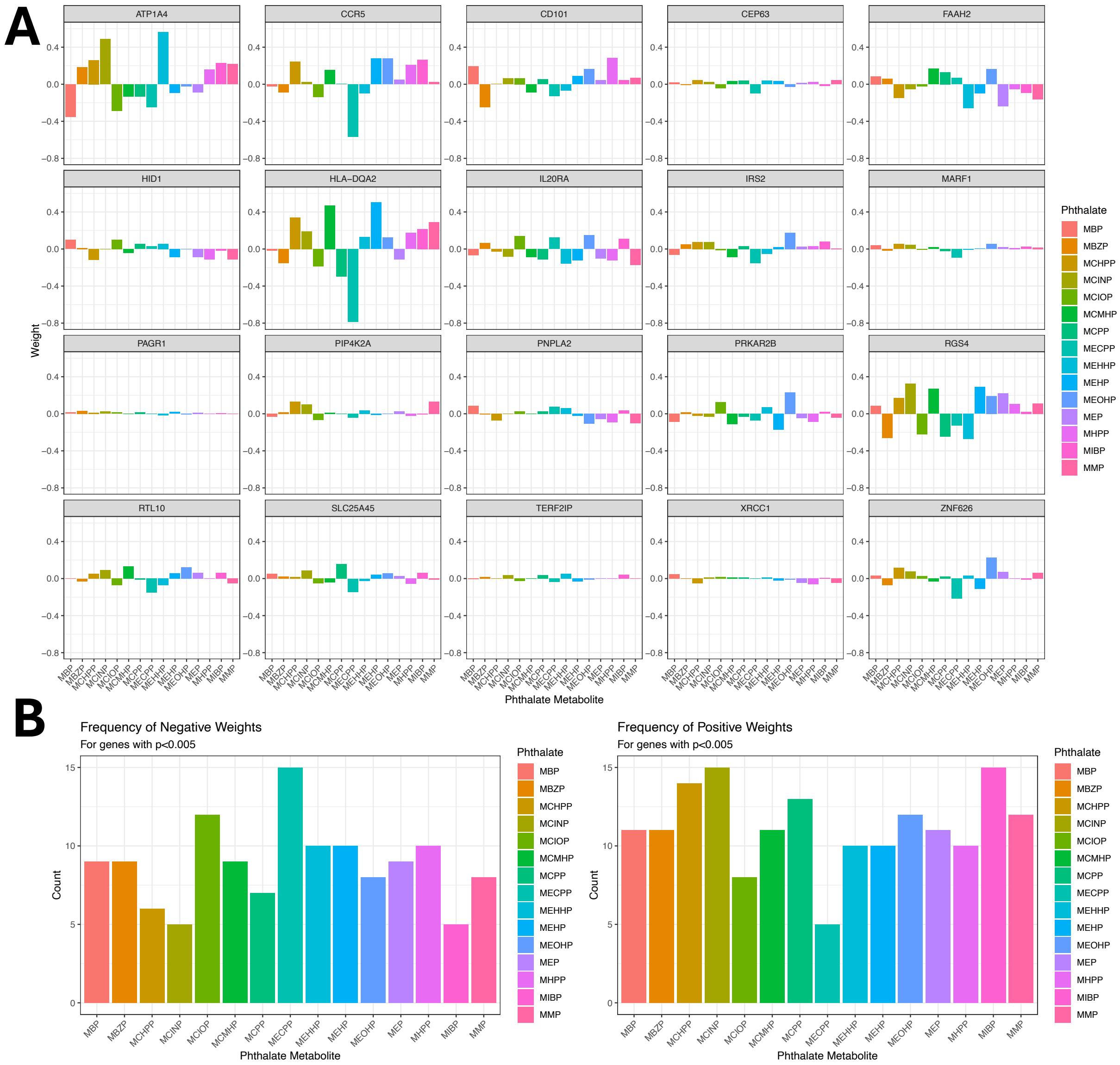


**Figure S6**: A) Contributions of phthalate mixture components to the overall mixture effect for each gene with p<0.005. B) Frequency of each mixture component being positively or negatively weighted for the 20 significant genes.

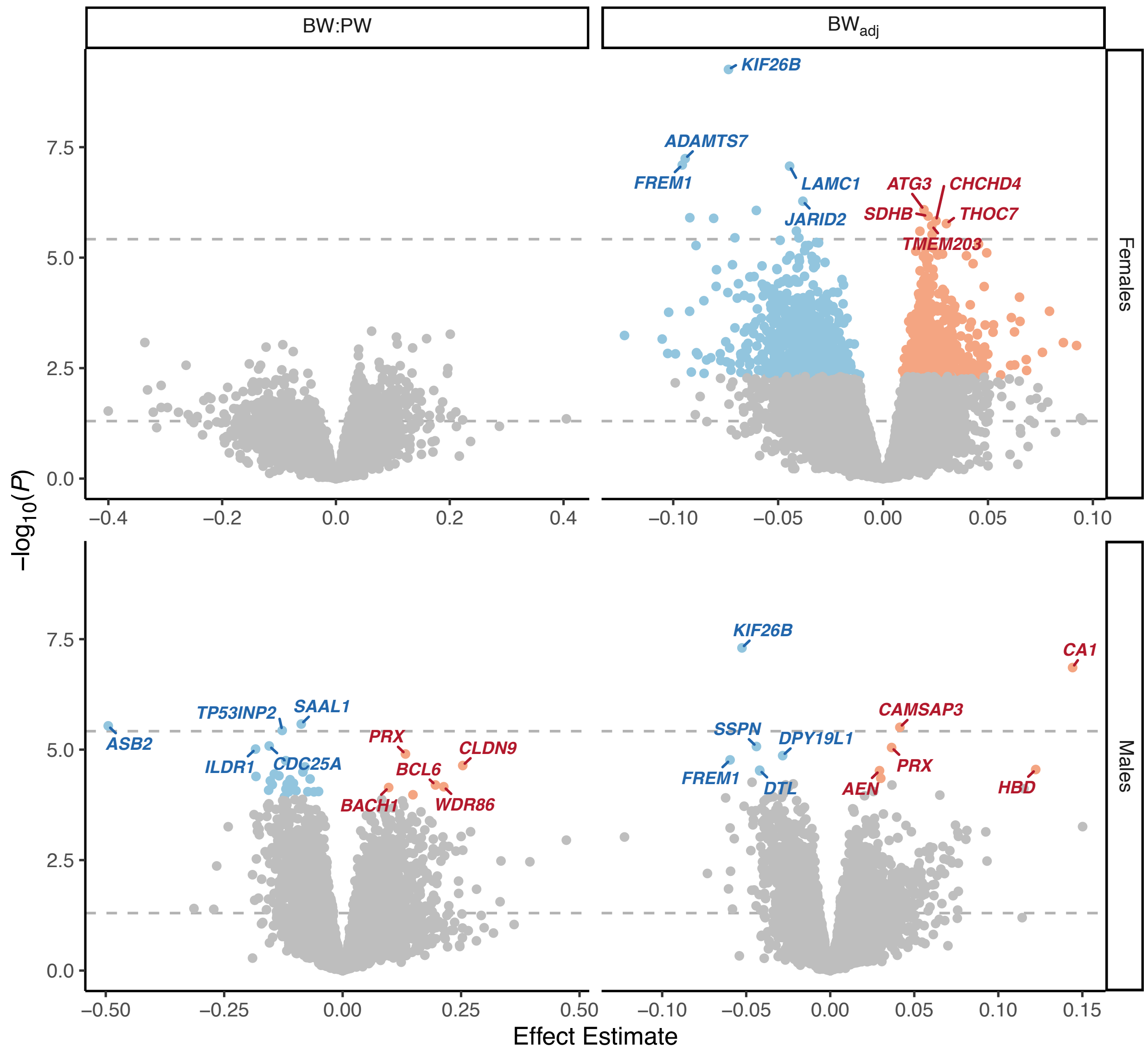


***Figure S7:*** Volcano Plots for sex-specific associations between placental gene expression and birthweight:placental weight ratio (BW:PW) and birthweight adjusted for placental weight ($BW_{adj}$). The top 5 up- and downregulated genes are labeled and all genes meeting FDR<0.05 are plotted in blue (downregulated) or red (upregulated). Dotted lines represent the Bonferroni and α=0.05 cutoffs.

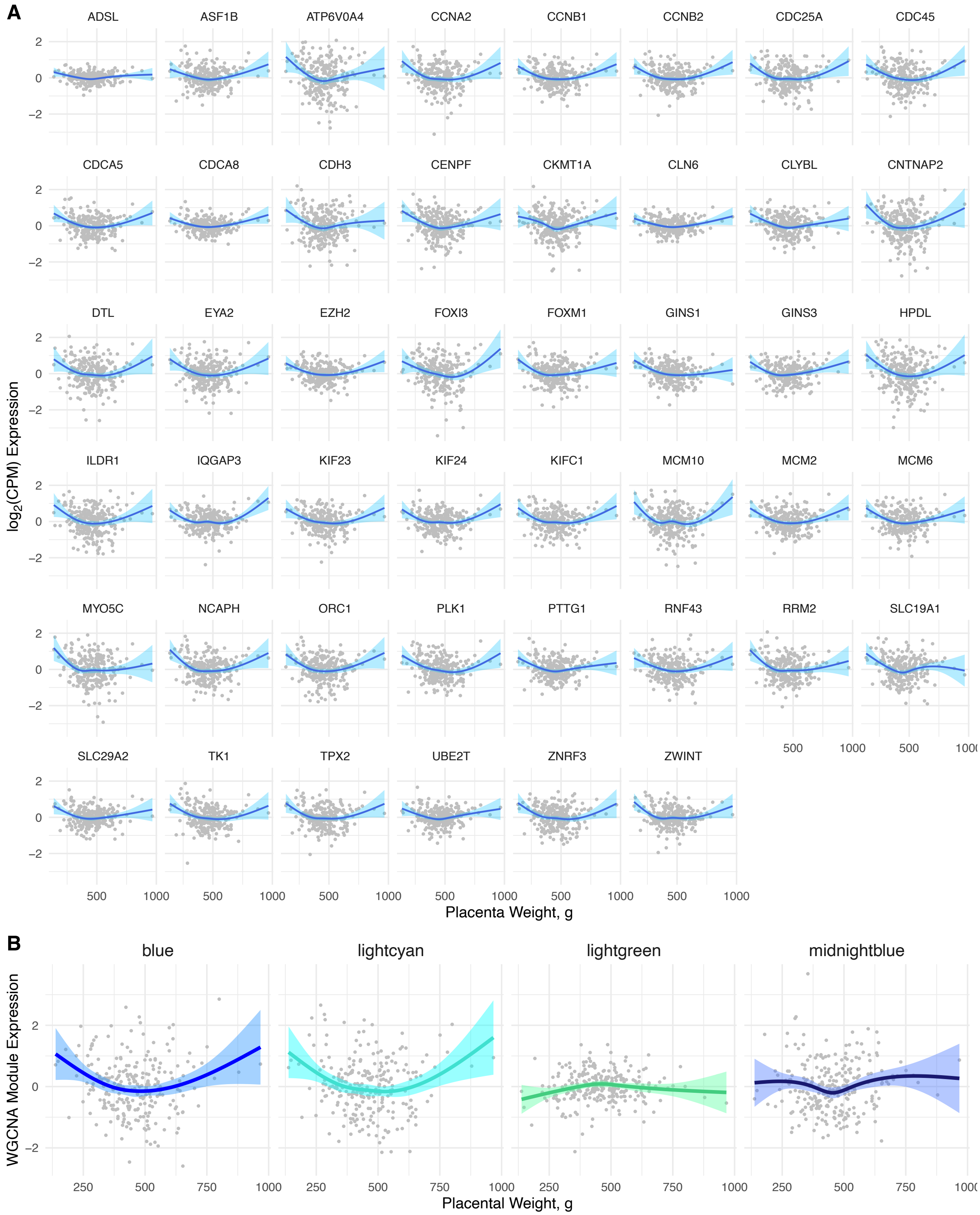


***Figure S8:*** Placental weight is nonlinearly associated with gene expression and coexpression. (A) 46 DEGs were significantly nonlinearly associated with placental weight (FDR<0.05). Gene expression is presented as covariate adjusted partial residuals (points) with the fitted natural cubic spline and 95% confidence interval. (B) 4 WGCNA modules were significantly nonlinearly associated with placental weight ($p<0.05$). Module expression is presented as covariate adjusted partial residuals (points) with the fitted natural cubic spline and 95% confidence interval.

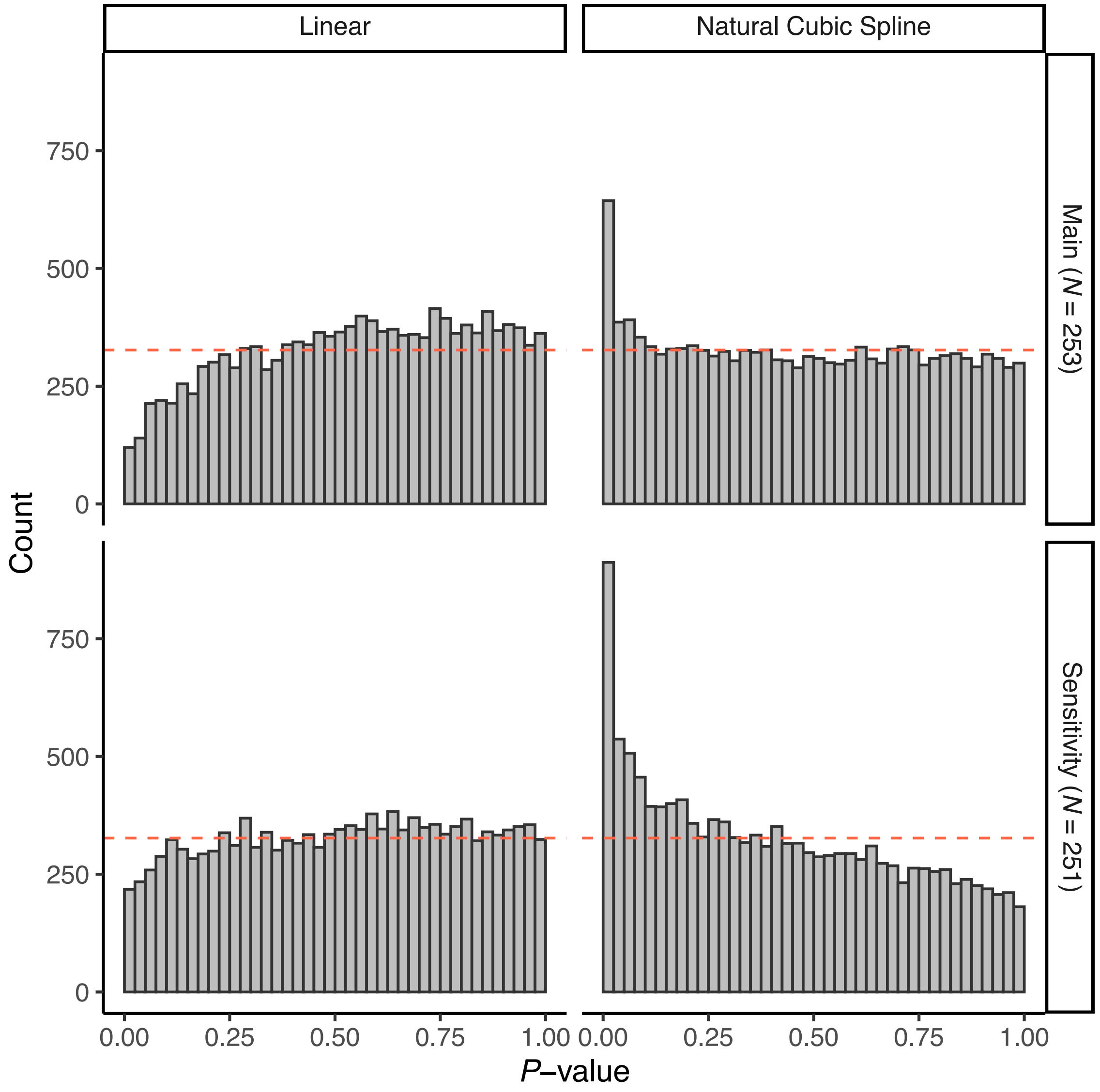


***Figure S9***: *P*-value distributions from the sensitivity analyses for the nonlinear association between placental weight and gene expression. Under the null hypothesis, the P-value distribution should be uniform. Modeling placental weight as a linear term (left column) suggested an improperly specified model in the main sample (top row, *N*=253) and sensitivity sample after removing the participants with the 2 highest placental weights (bottom row, *N*=251). Modeling placental weight as a nonlinear term (right column) resulted in more low values than expected by chance in the main sample (top row, *N*=253) and sensitivity sample (bottom row, *N*=251). The dashed line indicates the expected uniform distribution under the null hypothesis.

### SUPPLEMENTARY TABLES

***Table S1:*** Difference between entire PATHWAY GAPPS (PWG) sample and subset with RNA sequencing data.

| | Mediation Analysis (N=163) | RNA sequencing (*N*=465) | All PWG participants (*N*=673) | *P* value[1] |
|---|---|---|---|---|
| **Maternal Age (years)** | | | | 0.024 |
| Median (IQR) | 30 (26.5-34) | 31 (28-35) | 32 (28-36) | |
| Missing | 0 | 1 | 4 | |
| **Maternal BMI (kg/m²)** | | | | 0.425 |
| Median (IQR) | 25 (22-29) | 25 (22-30) | 25 (22-30) | |
| Missing | 0 | 55 | 74 | |
| **Gestational Age at Birth (weeks)** | | | | < 0.001 |
| Median (IQR) | 39.3 (38.3-40.2 | 39.1 (38.0-40.1) | 39.1 (37.6-40.1) | |
| Missing | 0 | 27 | 53 | |
| **Study Site** | | | | < 0.001 |
| Seattle | 56 (34.4%) | 212 (45.6%) | 345 (51.3%) | |
| Yakima | 107 (65.6 %) | 244 (54.4%) | 328 (48.7%) | |
| **Fetal Sex** | | | | 0.070 |
| Male | 83 (50.9%) | 252 (54.2%) | 349 (51.9%) | |
| Female | 80 (49.1%) | 213 (45.8%) | 324 (48.1%) | |
| **Labor status** | | | | 0.752 |
| Labor | 151 (92.6%) | 374 (88.8%) | 527 (88.6%) | |
| No labor | 12 (7.4%) | 47 (11.2%) | 68 (11.4%) | |
| Missing | 0 | 44 | 78 | |
| **Delivery Method** | | | | 0.097 |
| Vaginal | 115 (70.6%) | 297 (67.7%) | 410 (65.6%) | |
| C-section | 48 (29.4%) | 142 (32.3%) | 215 (34.4%) | |
| Missing | 0 | 26 | 48 | |
| **Maternal Smoking Status** | | | | 0.071 |
| No | 160 (98.2%) | 449 (97.4%) | 643 (96.5%) | |
| Yes | ≤5 | 12 (2.6%) | 23 (3.5%) | |
| Missing | 0 | 4 | 7 | |
| **Maternal Education** | | | | 0.359 |
| Less Than High School | 7 (4.3%) | 18 (3.9%) | 21 (3.1%) | |
| High School or GED | 50 (30.7%) | 122 (26.3%) | 181 (27.0%) | |
| Graduated Technical School or College | 70 (42.9%) | 204 (44.1%) | 292 (43.6%) | |
| Some Graduate work or Graduate Professional Degree | 36 (22.1%) | 119 (25.7%) | 176 (26.3%) | |
| Missing | 0 | 2 | 3 | |
| **Maternal Self-Reported Race** | | | | 0.281 |
| White | 128 (78.5%) | 358 (82.3%) | 518 (81.7%) | |
| Black/African American | ≤5 | 9 (2.1%) | 15 (2.4%) | |
| Asian | 6 (3.7%) | 24 (5.5%) | 32 (5.0%) | |
| Hawaii Native or Pacific Islander | ≤5 | ≤5 | ≤5 | |
| American Indian or Alaska Native | <5 | ≤5 | 12 (1.9%) | |
| Multiple | 17 (10.4%) | 28 (6.4%) | 40 (6.3%) | |
| Self-Reported Other | 6 (3.7%) | 11 (2.5%) | 16 (2.5%) | |
| Missing | 0 | 30 | 39 | |
| **Maternal Self-Reported Ethnicity** | | | | 0.003 |
| Not Hispanic | 139 (85.3%) | 383 (84.0%) | 568 (86.6%) | |
| Hispanic or Latino | 24 (14.7%) | 6739 (16.0%) | 88 (13.4%) | |
| Missing | 0 | 9 | 17 | |

1. Difference between RNA sequencing subset and full PWG cohort, evaluated by Kruskal-Wallis test (for continuous variables) or chi squared test (for categorical variables)

***Table S2:*** Summary of the concentration for 15 phthalate metabolites with >70% of measurements above the LOD. Each participant (N=222) had 1-3 measurements for each metabolite and concentrations were averaged by geometric mean to yield one value for each participant. DEHP is expressed as the molar sum (μM) of its monester metabolites (MEHP, MEHHP, MEOHP, MECPP, and MCMHP).

| **Metabolite** | **Parent Compound** | **Metabolite Molecular Weight** | **Median** | **Range** | **1st Trimester >LOD (N=51)** | **2nd Trimester >LOD (N=139)** | **3rd Trimester >LOD (N=189)** |
|---|---|---|---|---|---|---|---|
| | | *g/mol* | *ng/mL* | *ng/mL* | % | % | % |
| Monoethyl phthalate (MEP) | Diethyl phthalate (DEP) | 194.18 | 37.76 | 0.52-6594.3 | 100.00 | 99.28 | 100.00 |
| Mono (carboxyisooctyl) phthalate (MCIOP) | Diisononyl phthalate (DiNP); Di iso Decylphthalate (DiDP) | 322.40 | 17.79 | 0.90 - 824.39 | 100.00 | 100.00 | 99.47 |
| Monobutyl phthalate (MBP) | Dibutyl phthalate (DnBP); Butylbenzyl phthalate (BzBP) | 222.24 | 11.21 | 0.47 - 466.35 | 100.00 | 100.00 | 100.00 |
| Mono-2-ethyl-5-carboxypentyl phthalate (MECPP) | Di-2-ethylhexyl phthalate (DEHP) | 308.33 | 10.81 | 0.61 - 362.51 | 100.00 | 100.00 | 100.00 |
| Mono-2-ethyl-5-hydroxylhexyl phthalate (MEHHP) | Di-2-ethylhexyl phthalate (DEHP) | 294.35 | 9.40 | 0.52 - 2175.90 | 100.00 | 100.00 | 100.00 |
| Monobenzyl phthalate (MBZP) | Butylbenzyl phthalate (BzBP) | 256.25 | 8.24 | 0.14 - 283.55 | 96.08 | 99.28 | 99.47 |
| Mono[2-(carboxy-methyl) hexyl] phthalate (MCMHP) | Di-2-ethylhexyl phthalate (DEHP) | 308.33 | 8.003 | 0.29 - 230.98 | 100.00 | 100.00 | 100.00 |
| Mono-isobutyl phthalate (MiBP) | Diisobutyl phthalate (DiBP) | 222.24 | 6.32 | 0.04 - 163.00 | 100.00 | 100.00 | 98.94 |
| Mono-2-ethyl-5-oxo-hexyl phthalate (MEOHP) | Di-2-ethylhexyl phthalate (DEHP) | 292.33 | 5.33 | 0.08 - 1342.90 | 100.00 | 99.28 | 99.47 |
| Monocarboxy iso-nonyl phthalate (MCINP) | Diisodecyl phthalate (DiDP) | 336.40 | 2.30 | 0.19 - 64.25 | 100.00 | 100.00 | 100.00 |
| Mono-2-ethylhexyl phthalate (MEHP) | Di-2-ethylhexyl phthalate (DEHP) | 278.34 | 2.09 | 0.01 - 848.46 | 86.27 | 84.89 | 91.53 |
| Monomethyl phthalate (MMP) | Dimethyl phthalate (DMP) | 180.16 | 2.03 | 0.06 - 207.03 | 92.16 | 83.45 | 89.42 |
| Mono-3-carboxy-propyl phthalate (MCPP) | Dioctyl phthalate (DnOP); Di-n-butyl phthalate (DnBP) | 252.22 | 1.88 | 0.06 - 36.16 | 92.16 | 90.65 | 94.17 |
| Mono(4-hydroxypentyl) phthalate (MHPP) | Dipentyl phthalate (DPeP) | 252.26 | 0.55 | 0.01 - 139.22 | 98.04 | 94.96 | 94.71 |
| Mono-(7-carboxy-n-heptyl) phthalate (MCHPP) | Di-n-octyl phthalate (DNOP) | 308.33 | 0.31 | 0.02 - 14.89 | 94.12 | 84.89 | 94.18 |
| Di(2-ethylhexyl) phthalate (DEHP)* | - | 390.56 | 0.14 | 0.01-16.63 | - | - | - |
